# Antibodies against the capsid induced after intracranial AAV administration limits second administration in a dose dependent manner

**DOI:** 10.1101/2024.09.15.612566

**Authors:** Yuge Xu, Xiaoni Bai, Jianhua Lin, Kang Lu, Shihan Weng, Yiying Wu, Shanggong Liu, Houlin Li, Gong Chen, Wen Li

## Abstract

Recombinant adeno-associated virus (rAAV) is a widely used viral vector for gene therapy. However, a limitation of AAV-mediated gene therapy is that patients are typically dosed only once. In this study, we investigated the possiblility to deliver multiple rounds of AAV through intracerebral injections in the mouse brain. We discovered a dose-dependent modulation of the second round AAV infection by the first round AAV injection in the brain-wide scales besides the injection region. High-dose AAV infection increases chemokines CXCL9 and CXCL10 to recruit the parenchymal infiltration of lymphocytes. Surprisingly, the blood-brain-barrier was relatively intact. Brain-wide dissection discovered the likely rountes of the infiltrated lymphocytes through perivascular space and ventricles. Further analysis using B-cell depleted mice revealed that B lymphocytes, but not T lymphocytes, played a critical role in inhibiting the second round AAV infection. Strategies against neutralizing antibodies had limited effects, while reducing the dosage for the first injection or switching the second AAV to a different serotype appeared to be more effective in antagonizing the first round AAV inhibition. Together, these results suggest that mammalian brains are not immunoprivileged for AAV infection, but multiple rounds of AAV gene therapy are still possible if designed carefully with proper doses and serotypes.

## Introduction

Recombinant adeno-associated viruses (rAAVs) have been widely used as *in vivo* gene transfer vectors and clinical gene therapy because of moderate immunogenicity and their versatility in transferring genes of interest to a wide range of cell types.^1, 2, 3^ So far, there are at least five rAAV-based gene therapy products approved by the US Food and Drug Administration (FDA) (Luxturna for Leber’s congenital amaurosis, Zolgensma for spinal muscular atrophy, Hemgenix for hemophilia B, Elevidys for Duchenne muscular dystrophy, and Roctavain for hemophilia A), and 238 clinical trials employing rAAV-based gene therapy for various diseases.^3^ Ableit this great success, rAAV-based gene therapy still encounters many hurdles including immune responses and the loss of transgenes during long-term observation.^4, 5, 6^ Because AAV genomes mostly persist as episomes in transduced cells, the expression of transgenes may gradually be lost if the transduced target cells are dividing.^4, 5, 6^ In addition, like other viruses, rAAVs may trigger innate immune responses and adaptive immune responses.^4, 5, 6^ AAV-activated immune cells can negate the benefits of gene therapy. For example, T lymphocytes may kill the AAV-transduced cells serving as factories for producing therapeutic proteins. Therefore, re-administration of rAAV vectors may be a desirable approach to achieve long-term therapeutic effects.^4, 5, 6^

The major challenge in rAAV vector re-administration is the formation of neutralizing antibodies (NAbs) following the first round of AAV infection, which has been widely investigated when targeting cells in the peripheral system (e.g. liver and muscle)^6, 7^. However, studies on the ability to readminister rAAV in the brains are limited and controversial. Some studies reveal that repeat administration of rAAV in the brain is both serotype- and transgene-dependent.^8, 9, 10^ Moreover, immunosuppression after the first round AAV administration was considered transient and to be triggered beyond generating NAbs.^9, 10^ These findings are reasonable based on the previous perspective that central nervous system is “immune-privileged” due to the structure of blood-brain-barrier (BBB) that restricts the invasion of immune cells.^11^ As the understanding of brain immunity evolves with the identification of specialized immunological compartments within the brain, including the meninges, the choroid plexus and the perivascular spaces, immune cells at the borders of the brain may provide surveillance and modulate the development and functions of neuronal circuits.^11, 12, 13^ Thus, the immune response induced after the intracranial administration of AAV and its effect on the multiple rounds of AAV gene therapy remains to be further explored.

In this study, we examined re-administration of rAAV directly into the brain parenchyma and discovered the compromising effect of the first round of rAAV in a dose-dependent manner. High-dose of first round rAAV suppressed the transgene expression of the second round rAAV of the same serotype. The compromising effect extended brain wide and sustained at least 5 months after the first injection. Using bulk RNA sequencing (RNA-seq) analysis, immunostaining and *in vitro* NAb assay, we discovered the invasion of B lymphocytes and the production of NAbs in both injection site and non-injection regions. Surprisingly, the infiltration of B cells was not due to the BBB breakdown induced by high-dose AAV intracranial injection. Instead, they invaded through the perivascular spaces and ventricles in the injection regions as a result of the increased permeability of brain-cerebrospinal fluid barrier (BCSFB) along the lateral ventricles, and recruited by chemockines CXCL10 and CXCL9 in the brain parenchyma. The immunosuppression of the first-round AAV failed to be circumvented through the systemic administration of IgG-degrading enzyme or AAV empty vectors against neutralizing antibodies. But reducing the dosage of the first round AAV injection or switching the second round AAV to a different serotype appeared to be more effective to antagonize the first round AAV inhibition. Together, this study illuminates the antiviral response from the border immune niches of CNS, and provide potential solutions for intracerebral re-administration of rAAVs.

## Results

### Administration of high-dose AAV compromises the transduction of second round AAV of the same serotype

To explore the possibility of multiple rounds of gene transduction in the brain through intraparenchymal injection of AAV, we investigated the effect of first round AAV injection on the second round AAV injection by exploring a variety of parameters such as serotypes, dosage, and promoters. AAV5 expressing tdTomato (red) under the astrocyte-specific promoter GFAP104 was injected into the striatum for the first round of infection,^14, 15^ and after a 30-day interval, a second round of AAV5 expressing GFP (green) under the same GFAP104 promoter was administrated at the same position (Fig. 1A). We tested three different dosages, a high dose of 4E10 vg per mouse, a middle dose of 1.2E10 vg per mouse and a low dose of 2E9 vg per mouse in the first round of injection to explore the dosage effect (Fig. 1A). The expression levels of GFP at 14 days after the second injection were examined. As a control, mice that received PBS as the first round of injection and GFP-expressing virus as the second round of injection showed wide-spreading and high expression level of GFP (Fig. 1B). The GFP expression representing the second round of virus transduction after the injection of the low-dose tdTomato-expressing virus as the first injection seemed normal to that receiving PBS as the first injection (Fig. 1C). In addition, 1^st^ and 2^nd^ round of virus can be transduced in the same cells in the low-dose group (indicated by cells double positive for tdTomato and GFP, Fig. S1A). However, in the mice that received middle- and high-dose first round injection of the tdTomato-expressing virus, the expression of second round injection of GFP was completely undetectable (Fig. 1D, Fig. S1A-B). Quantitatively, both the expression area and the intensity of GFP were comparable between mice receiving PBS and low dose of the tdTomato-expressing virus as the first injection, which was barely detectable in the mice receiving the middle- and high-dose AAV5 tdTomato as the first injection (Fig. 1H and I). This result indicates that intrastriatal administration of high-dose AAV5 compromises the transgene expression transduced by the second-round AAV5. Previous study has reported that delaying the re-administration to an 11-week interval abrogated the immune response in the second injection site.^9^ However, we found that extending the interval between the two injections to even 150 days had no improvement in the GFP expression level of the second round GFP injection, suggesting that the high dose AAV5 has a long-term compromising effect (Fig. 1E-G,1J). Changing the order of transgene expression (AAV5 GFAP104::GFP as the first injection and AAV5 GFAP104::tdTomato as the second injection, Fig. S1C, S1D, and S1I), or increasing the dose of the virus of the second round (from 2E9 vg/mouse to 4E10 vg/mouse, Fig. S1E, S1F, and S1J) failed to circumvent the compromising effect of the first round high-dose AAV with the same serotype. In fact, this compromising effect of high-dose AAV on gene transduction of the second round of AAV is not limited to AAV5 but can be applied to AAV9 as well. The administration of high-dose AAV9 GFAP104::tdTomato greatly compromised the second round AAV9 GFAP104::GFP expression (Fig. S1G, S1H, and S1K). Moreover, the compromising effect of the high-dose AAV is also independent of the transduced cells that express transgenes. Changing the promoter from GFAP104 to synapsin promoter that targets neurons, we also observed that the high-dose AAV9 Syn::mCherry significantly compromised the expression of second round AAV9 Syn::GFP in neurons (Fig. 1K-N). Even more interestingly, transducing astrocytes with high-dose AAV5 GFAP::tdTomato as the first round injection compromised the expression of second round AAV5 SYN::GFP in neuronal cells, and vice versa (Fig. S2). Taken together, these results demonstrate that when conducting multiple rounds of intracerebral injection of AAV, it is critical to select the right dose for the first round of injection, because the first round of high-dose AAV may severely compromise the transduction of the second round AAV of the same serotype.

**Figure 1.**
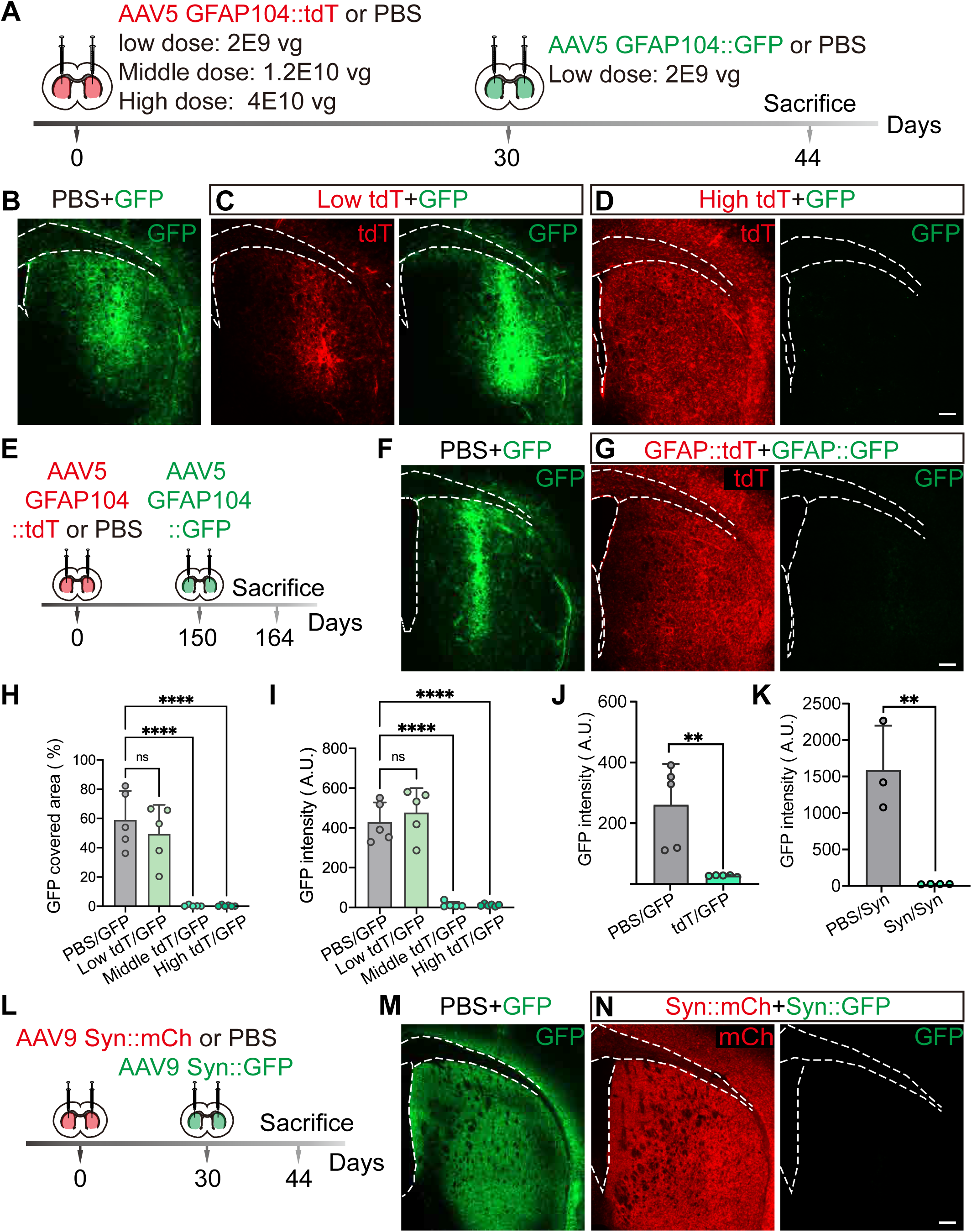
Intrastriatal administration of AAV compromises the transduction of second round AAV of the same serotype in a dosage manner. (A) Experimental design: intrastriatal delivery of PBS (as the control) or different dosages of AAV5 GFAP104::tdTomato (tdT) followed by second-round injection of AAV5 GFAP104:: GFP 30 days later. vg, viral genome. (B-D) Representative images showing the expression of second round of AAV5-GFP (green in B, C and D) and first round of AAV5-tdT (red in C and D) in PBS (B), low-dose (2E9 vg/mouse, C), or high-dose (4E10 vg/mouse, D) group. Note that the expression of second round of AAV-GFP is completely abolished in the high-dose group. Scale bar: 200 μm (E) Experimental design: re-administration of high-dose AAV5 GFAP104::GFP 150 days after first round injection of AAV5 GFAP104:: tdTomato (tdT). (F-G) Representative images showing the expression of second round of AAV5-GFP (green in F and G) and first round of AAV5-tdT (red in G) in PBS (F) and high-dose (G) group at a 150-day interval. Note that the expression of second-round AAV-GFP is completely abolished in the mice receiving high-dose AAV5 in the first round. Scale bar: 200 μm (H-I) Quantitative analyses showing a significant reduction in the infection area (H) and intensity (I) of second-round AAV5 following the first-round injection of high- and middle-dose AAV5, but not PBS or low-dose AAV5. Mean (SD); one-way ANOVA with Dunnett’s multiple comparisons, ****P< 0.0001, n= 5 mice/group. (J) Quantitative analyses showing a dramatic inhibition of the second round GFP intensity by intrastriatal delivery of high-dosage AAV5 at a 150-day interval. Mean (SD); unpaired t-test, **P= 0.0047, n= 5 mice/group. (K-N) Experimental design to investigate the compromising effect of high-dose AAV9 using Syn promoter on the following AAV9 redosing (L). The representative images (M and N) and quantitative analyses (K) revealing that GFP expression indicated the second round of AAV9 is compromised by the first injection of high dose AAV9, but not PBS. Mean (SD); unpaired t-test, ****P< 0.0001, **P= 0.0047 for J, **P= 0.0032 for K, n= 3 mice/group; scale bars: 200 μm.

### Time course and brain-wide compromising effects of high-dose AAV

We next investigated when this compromising effect occurred by testing a variety of time intervals between the first and second round of AAV injections. Mice that received the high dose of AAV5 GFAP104::tdTomato (4E10 vg) as the first intrastriatal injection were administrated AAV5 GFAP104::GFP (2E9 vg) as the second injection at different time intervals (7-, 14-, 21- and 30- day interval). Intrastraital injection of the two-round viruses with a 7-day interval caused a moderate decrease in the second round GFP expression (Fig. 2A). Attenuation of the second round GFP expression became significant after a 14-day interval, and almost undetectable at intervals of 21 and 30 days (Fig. 2A). Quantitative analyses revealed that the expression of the first round of virus was not affected by the second round of virus injection at any intervals (Fig. 2B), but the second round of AAV expression was severely compromised by the first round of high-dose AAV (Fig. 2C). These results suggest that the compromising effect of high-dose AAV is time dependent, taking place around 7 to 14 days after the first injection.

**Figure 2.**
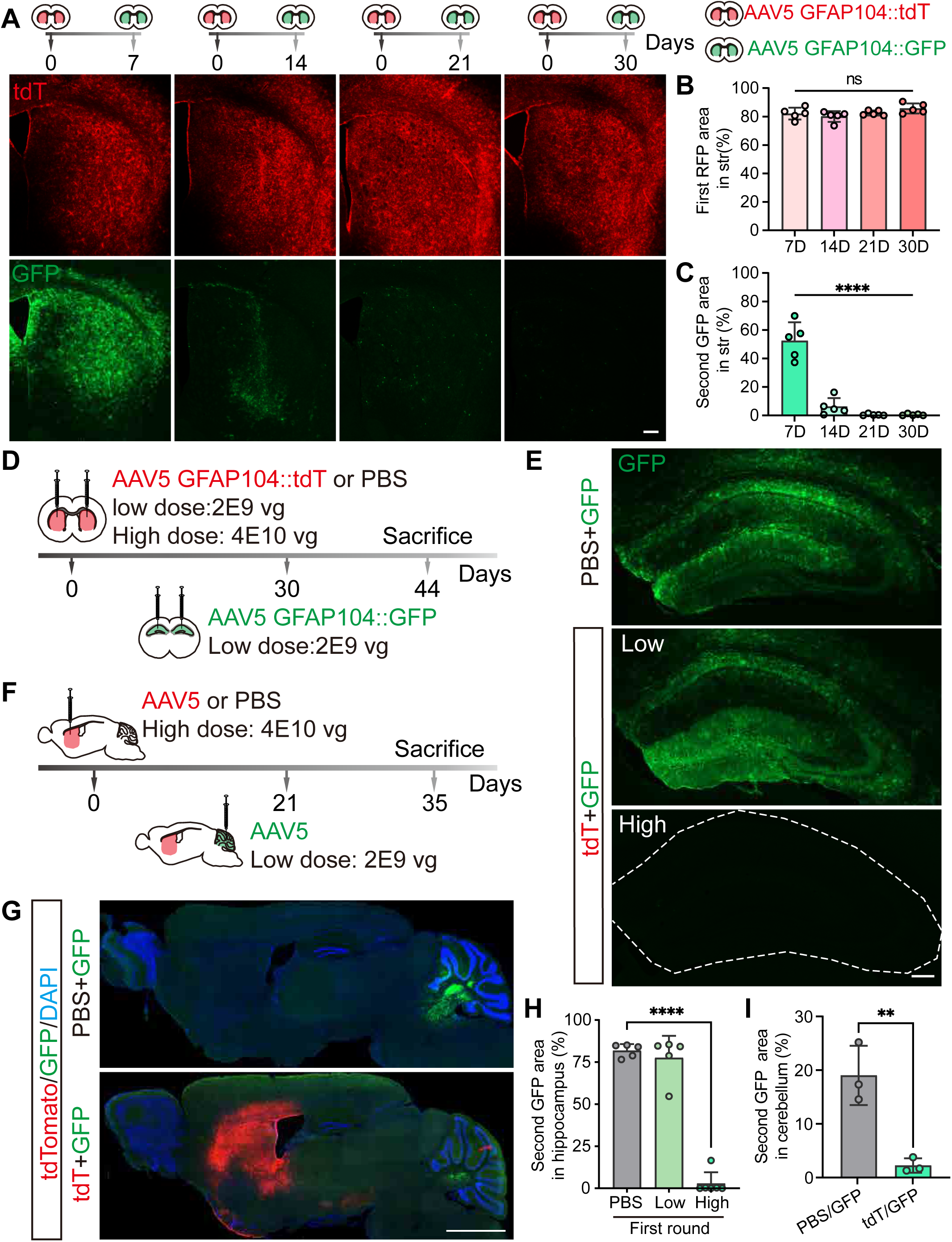
The compromising effects of high-dose AAV is time-dependent and in brain wide-scaled. (A) Representative images revealing that first round high-dose AAV5 (red) compromised the expression of second round AAV5 (green) in a time-dependent manner. Scale bar: 200 μm (B-C) Quantitation of the infection area of first-round AAV5 (B) and second-round AAV5 (C) with different time intervals. Mean (SD); one-way ANOVA with Dunnett’s multiple comparisons, ****P< 0.0001, n= 5 mice/group. (D) Experimental design to investigate the compromising effects in the hippocampus as a non-injection region adjacent to the striatum receiving the first round of AAV microinjection. PBS is used as the control. (E) Representative images showing that intrastriatal injection of high-dose AAV compromises the expression of second round AAV in the hippocampus. The hippocampus is delineated by the white dashed line. Scale bar: 200 μm (F) Experimental design to investigate the compromising effects in the cerebellum as a non-injection region distant from the striatum receiving the first round of AAV microinjection. PBS is used as the control. (G) Representative images showing that intrastriatal injection of high-dose AAV suppresses the GFP expression in the cerebellum. The cell nuclei are counterstained with DAPI (blue). Scale bar: 1000 μm (H-I) Quantitation of the GFP expression of the second-round AAV5 in the hippocampus (H) and cerebellum (I). Note that the high-dose AAV5 administered in the striatum inhibits the expression of second round of AAV5 delivered in brain regions not receiving the first round AAV injection. Mean (SD); one-way ANOVA with Dunnett’s multiple comparisons or unpaired t test, ****P< 0.0001, **P= 0.0069, n= 3-5 mice/group.

After examining the time course, we next asked whether the compromising effect was restricted to the injection site or extended to other brain regions. We first changed the second injection site to the hippocampus adjacent to the striatum (Fig. 2D), and found that the second round of GFP expression in the hippocampus was severely compromised by high, but not low, dose of the first-round viral infection in the striatum (Fig. 2E and quantified in 2H). We then changed the second injection site to the more distant cerebellum (Fig. 2F), and discovered that the second round of GFP expression in the cerebellum was also dramatically compromised by high-dose AAV infection in the striatum (Fig. 2G). However, different from the nearly complete suppression of GFP expression in the hippocampus, the high-dose of intrastriatal injection of AAV5 allowed weak expression of GFP in the cerebellum as the second round of viral injection (Fig. 2G and quantified in 2I). These results suggest that the compromising effect of high-dose AAV may occur in wide brain regions away from the injection sites.

### High-dose AAV induces immune response in both injection site and non-injection site

To investigate the molecular mechanisms of the compromising effects of high-dose AAV transduction, we performed bulk RNA-seq to examine the global gene expression changes in both injection site (striatum) and non-injection site (hippocampus) following high-dose AAV5 GFAP104:tdTomato injection (Fig. S3A). We identified 222 differentially expressed genes (DEGs, FDR < 0.05) in the striatum of mice at 21 days following high-dose AAV5 injection compared with that of PBS control, with 189 upregulated genes and 33 downregulated genes (Fig. 3A). In contrast, the number of DEGs induced in the non-injection site, the hippocampus, was 58, only one-third of that in the striatum (Fig. S3B). Kyoto Encyclopedia of Genes and Genomes (KEGG) analysis revealed that many DEGs in both striatum (Fig. 3B) and hippocampus (Fig. S3C) were enriched in immune system pathways, including infectious diseases (e.g. Epstein-Barr virus, COVID-19, etc.), immune responses (e.g. antigen processing and presentation, phagosome, etc.) and immune diseases (e.g. autoimmune thyroid disease, viral myocarditis, etc.). Since most of the DEGs were upregulated genes (189/222 in striatum, 54/58 in hippocampus, Fig. 3A and S3B), further analyses were mainly focused on these upregulated DEGs. The volcano plots showed that the striatum and hippocampus shared a lot of transcripts enhanced by high-dose AAV5 treatment (eg. *Igkc, H2-K1, Irgm2, etc.*), although the number of the upregulated transcripts in the striatum was much more than that in the hippocampus (Fig. S3D and S3E). This result implied that high-dose AAV5 induced common changes in both injection site and non-injection region. To further illuminate these common changes, we compared the enhanced transcripts in these two regions and found 51 overlapped transcripts (Fig. 3C). The heatmap revealed the top 30 of the shared up-regulated transcripts in the striatum and hippocampus after the injection of high-dose AAV, which were associated with immunoglobin domains (*Igkc*, *Jchain*, *Ighm*, *Ighg2b*), antigen presentation (*H2-D1*, *H2-K1*, *H2-Aa*, *H2-Q7*), and anti-viral response (*Irf7*, *Irgm1*, *Igrm2*, *Ifit3*) (Fig. 3D and 3E).^16, 17, 18, 19^ The shared up-regulated transcripts displayed higher expression level in the striatum than in the hippocampus (Fig. 3D). These results discovered that the administration of high-dose AAV5 induced common immune responses including immunoglobin protein synthesis, antigen presentation, and anti-viral response in both injection and non-injection regions.

**Figure 3.**
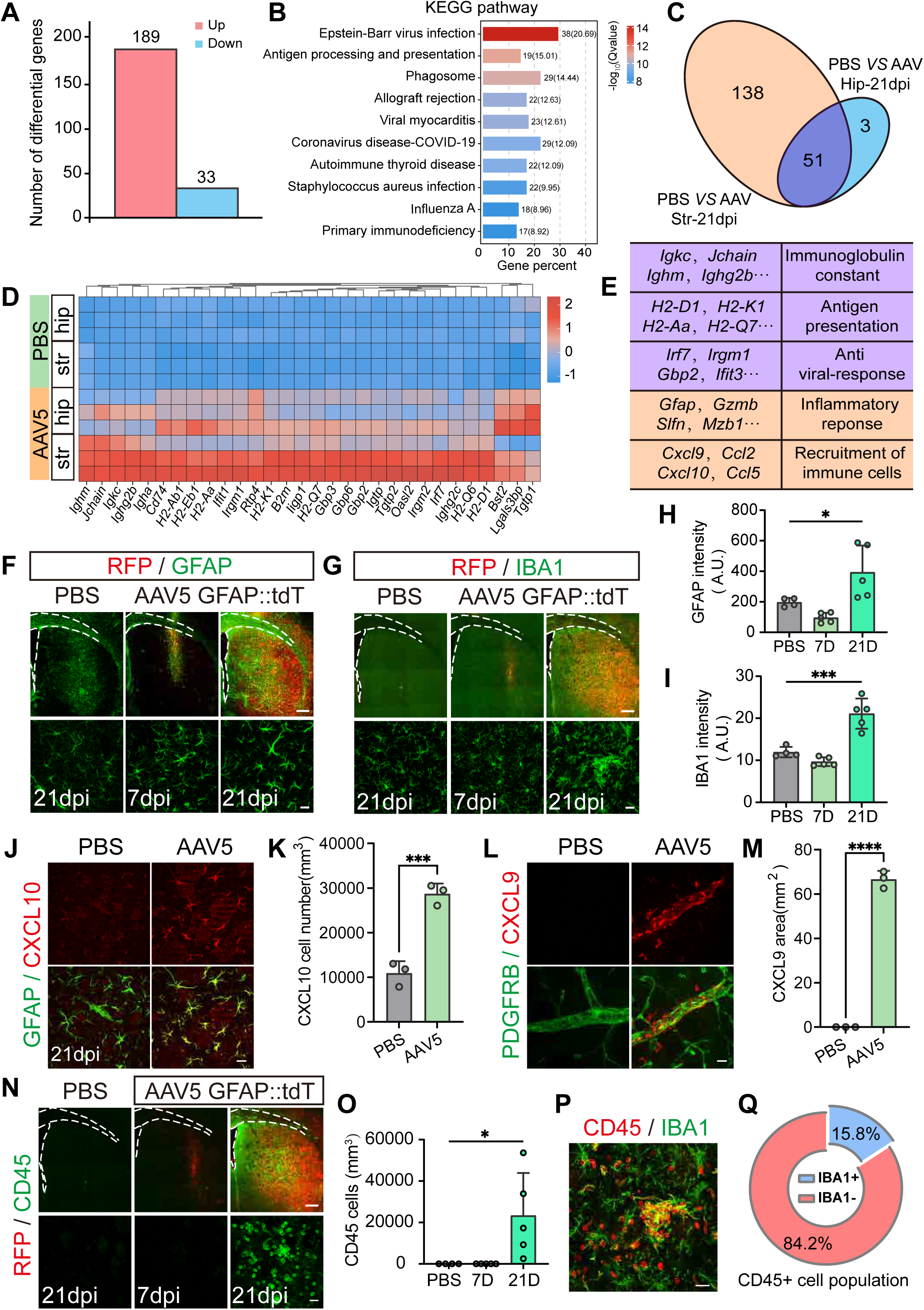
High-dose AAV induces immune response in both injection site and non-injection site. (A) Histogram showing 222 differentially expressed genes (DEGs, FDR< 0.05) in the striatums between the high-dose AAV5-treated mice and PBS-treated mice (n= 3 mice/group). (B) The KEGG-enriched bar graph showing that the pathways in the striatum induced by intrastriatal microinjection of high-dose AAV are mainly restricted to the immune response (one-way ANOVA analysis with Tukey test). (C) Venn Diagram showing that there are 51 upregulated DEGs shared between the striatum and hippocampus after high-dose AAV5 intrastriatal administration, while there are 138 and 3 upregulated DEGs unique to the striatum and hippocampus, respectively. (D) Heatmap showing the top 30 of upregulated DEGs shared between the striatum and hippocampus after high-dose AAV injection. (E) The table showing three classes of DEGs shared between the striatum and hippocampus and two classes of DEGs unique to the striatum. (F-G) Immunostaining of GFAP (green, F) and IBA1(green, G) in the injected areas (RFP in red). The corpus callosum and lateral ventricle are delineated by the white dashed lines. Scale bars: 200 μm for low-magnification images and 20 μm for high-magnification images; dpi: days post injection (H-I) Quantification of the intensity of GFAP (I) and IBA1 (J) covered area within the injected sites. Mean (SD); one-way ANOVA with Dunnett’s multiple comparisons, *P= 0.0361, ***P= 0.0002, n= 5 mice/group. (J-K) Representative images (J) and quantitation (K) showing the elevated expression of chemokine CXCL10 (red) in reactive astrocytes (GFAP, green) induced by high-dose AAV5. Scale bars: 20 μm; dpi: days post injection. Mean (SD); unpaired t-test, ***P= 0.0010, n = 3 mice/group. (L-M) Representative images (L) and quantitation (M) showing that high-dose AAV5 increase the chemokine ligand CXCL9 (red) around blood vessels (PDGFRB, green). Scale bars: 20 μm. Mean (SD); unpaired t-test, ****P< 0.0001, n= 3 mice/group (N-O) Representative images (N) and quantitation (O) revealing the massive invasion of CD45^+^ cells (green) in the injected areas (RFP in red). The corpus callosum and lateral ventricle are delineated by the white dashed lines. Scale bars: 200 μm for low-magnification images and 20 μm for high-magnification images; dpi: days post injection. Mean (SD); one-way ANOVA with Dunnett’s multiple comparisons, *P= 0.0309, n= 5 mice/group. (P-O) Representative images (P) and quantitation (O) showing that more than 80% CD45 positive cells (red) were IBA1 negative (green) in the AAV injected areas at 21 days. Scale bar: 20 μm.

Besides the common changes, there were 138 transcripts unique to the striatum enhanced by high-dose AAV5 treatment (Fig. 3C). These up-regulated transcripts were related to inflammatory response (*Gfap*, *Gzmb*, *Slfn*, *Mzb1*) and recruitment of immune cells (*Cxcl9*, *Ccl12*, *Cxcl10*, *Ccl5*) (Fig. 3E),^20, 21, 22, 23, 24, 25, 26^ suggesting a broad activation of local inflammation and immune cell chemotaxis in the region where the virus was injected. Consistent with this, we observed robust activation of astrocytes (indicated by elevated signal of GFAP and VIMENTIN, Fig. 3F, 3H, Fig. S3F And S3G) and microglia (indicated by elevated signal of IBA1 and CD68, Fig. 3G, 3I, S3H, and S3I) in the striatal region 21 days after the injection of high-dose AAV.^25, 27, 28^ Interestingly, the reactive astrocytes induced by high-dose AAV5 showed a high expression level of CXCL10 (Fig. 3J, 3K, and S3J), a chemokine ligand to recruit immune cells^20, 29, 30^. In addition, CXCL9, another chemokine ligand responsible for immune cell recruitment,^20, 29^ was observed expressed in some of astrocytes in the striatum at 21 dpi following high-dose AAV injection (Fig. S3L). More frequently, it was observed in cells around the blood vessels and between the pericyte layers and astrocytic limitan (indicated by PDGFRB and GFAP) (Fig. 3L, 3M, S3K and S3L).^31, 32^ In accordance with the increase of CXCL10 and CXCL9, there were massive CD45^+^ cells in the striatum 21 days after the injection of high-dose AAV5 (Fig. 3N and 3O). More than 80% of the CD45^+^ cells were IBA1 negative, suggesting that they were monocytes infiltrated into the parenchyma rather than local microglia (Fig. 3P and 3Q).^33, 34^ In constrast, in the hippocampus where the virus was not directly injected, the activation of local glia, the elevation of chemokines (CXCL10 and CXCL9) and infiltration of leukocytes was mild (Fig. S3M). These data suggested that high-dose AAV5 induced the activation of chemokines CXCL9 and CXCL10 mainly in the injection sites, which in turn recruited leukocytes into the brain.

### High-dose AAV triggers B cell infiltration and production of neutralizing antibodies to compromise the re-administration of AAV

The above transcriptomic analysis revealed an upregulation of transcripts associated with immunoglobulin domains following high dose AAV injection, which prompted us to examine potential AAV neutralizing antibodies (NAbs) and B cells that produce NAbs. B cells are barely detectable in the brain under normal conditions.^35, 36^ But 21 days after the high-dose AAV5 injection, immunostaining identified both B220^+^ B cells and CD138^+^ plasma B cells in the striatum, while they were absent in the striatum receiving PBS or at early stage after viral injection (Fig. 4A-C). Consistently, the number of IgG^+^ cells as well as the signal of IgG in the striatum increased dramatically at 21 days after high-dose AAV5 injection (Fig. 4A). Quantitively, the intensity of IgG elevated 9 times in the striatum 21 days after the viral injection (Fig. 4D). The infiltration of B cells was also observed in the hippocampus (Fig. S4A). These data suggested that high-dose AAV triggered the infiltration of antibody-secreting cells (ASCs) into both the injection and non-injection sites.

**Figure 4.**
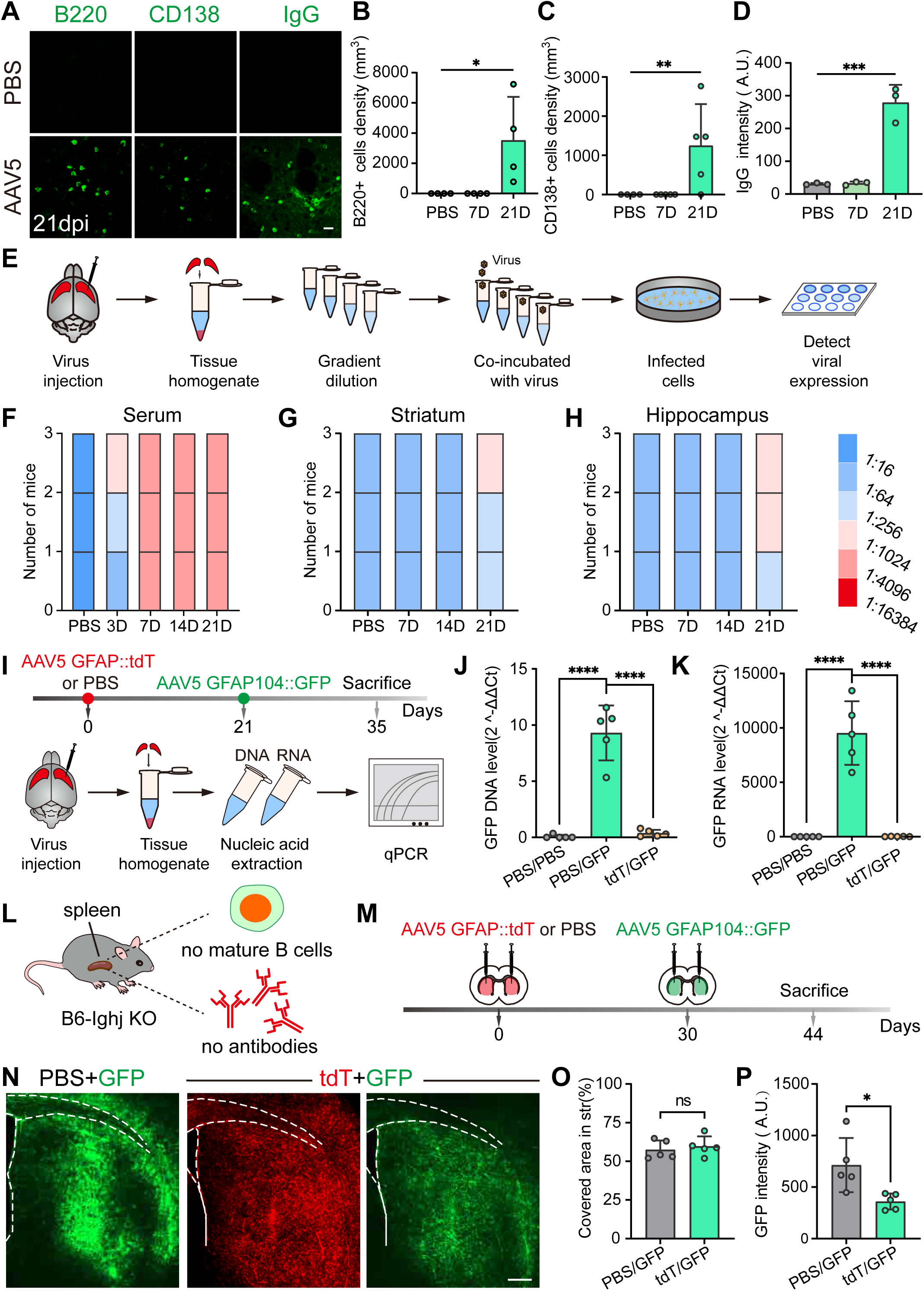
High-dose AAV triggers B cell infiltration and production of neutralizing antibodies to compromise the re-administration of AAV. (A) Representative images showing the invasion of B220^+^ (labeling pan B cells, left), CD138^+^ (labeling plasma cells, middle), and IgG^+^ (labeling immunoglobulin, right) cells in the striatum 21 days after receiving high-dose AAV5. Scale bar: 20 μm; dpi: days post injection. (B-C) Quantitative analysis showing that the density of B220^+^ (B) and CD138^+^ cells (C) increases in the striatum receiving high-dose AAV5. Mean (SD); one-way ANOVA with Dunnett’s multiple comparisons, *P= 0.0277 for B and **P= 0.0027 for C, n= 3 or 4 mice/group. (D) Quantitative analysis showing that the intensity of IgG increases in the striatum receiving high-dose AAV5. Mean (SD); one-way ANOVA with Dunnett’s multiple comparisons, ***P= 0.001, n= 3 or 4 mice/group. (E) Experimental procedure of in vitro neutralizing antibodies (NAbs) assay. (F-H) Stacked histogram showing the titer of neutralizing antibody in the serum (F), striatum (G) and hippocampus (H) collected at different time points after PBS/AAV5 injection. Samples are categorized by the AAV NAb titer that is presented on a colored scale. N = 3 mice (I) Experimental design to investigate the infection of 2nd round of AAV5 in the striatum: all mice were injected PBS/AAV5 GFAP104::tdTomato and AAV5 GFAP104::GFP with a 21-day interval. Two weeks later, the mice were sacrificed for qPCR analysis. (J-K) The qPCR analysis showing that the second transgene (GFP) is hardly detectable at either DNA (J) or RNA (K) level in mice that received the first high-dose AAV5 injection. Mice that received two rounds of PBS serve as negative controls. Mean (SD); one-way ANOVA with Dunnett’s multiple comparisons, ****P< 0.0001, n= 5 mice/group. (L) Schematic diagram showing the lack of mature B cells and antibodies in B6-Ighj-KO mice. (M) Experimental workflow: B6-Ighj KO mice receiving either PBS or high-dose AAV5 GFAP104::tdTomato in the striatum as the first injection received AAV5 GFAP104::GFP injection at a 21-day interval. 14 days after the second round of injection, the mice were sacrifice for analysis. (N) Representative images showing that the GFP expression level in the mice receiving AAV5 expressing tdTomato as the first injection was comparable to that of mice received PBS. The corpus callosum and lateral ventricle are delineated by the white dashed lines. Scale bar: 200 μm (O-P) Quantified data showing the similar GFP coverage (O) and intensity (P) in PBS and AAV5-treated B6-Ighj KO mice. Mean (SD); unpaired t test, *P= 0.0250, n= 5 mice/group.

The presence of B cells and IgG^+^ cells suggested a potential inhibition of 2^nd^ round AAV infection by NAbs following the 1^st^ round high-dose AAV injection. To test this possibility, we collected the sera and parenchymal tissues of the striatum and hippocampus at different time points (7 days, 14 days, and 21 days) after injecting AAV5 GFAP104::tdTomato into the striatum, and performed *in vitro* NAb assay (Fig. 4E). After the intrastriatal injection of high-dose AAV5, we detected NAbs in the sera as early as 3 dpi and the titer increased rapidly to a high level (over 1:4096) at 7 dpi (Fig. 4F). The detection of NAb in the striatum and hippocampus was relatively later, which was detected until 21 days after the injection (Fig. 4G and 4H). The titer of the parenchymal NAb was lower (titer < 1:1024) than that of sera NAb, but it was sufficient to inhibit the redose of the AAV. As shown by the real-time qPCR, both GFP DNA and RNA transcripts (indicators of second round of AAV5 GFAP104::GFP) were barely detectable in the striatum of mice that received the first round of high-dose AAV5 GFAP104::tdTomato, whereas both GFP DNA and RNA transcripts were detected in mice receiving the first-round injection of PBS (Fig. 4I-K). To further verify that it is the B cells and the production of NAbs to inhibit the redose, we employed the Ighj knockout mice (B6-Ighj KO) that are deficient in developing B cells to perform the redosing experiment (Fig. 4L).^37^ As reported,^37^ the spleens of Ighj knockout mice were smaller than those of wild-type mice (Fig. S5B), and were absent of mature B cells (indicated by B220^+^) in the germinal center of the spleens (Fig. S5C). When GFP-expressing AAV5 was injected in the striatum of B6-Ighj KO mice 30 days after receiving the first round of injection of high-dose tdTomato-expressing AAV5 (Fig. 4M), the GFP signal was clearly detected in broad areas of the striatum (Fig. 4N-P), suggesting that reducing B cells relieved the compromising effect of high-dose AAV.

In addition to the B cells, we also discovered massive infiltration of T cells in the brain tissues after the high-dose AAV5 injection, including CD8^+^ cytotoxic T lymphocytes (CTL) and CD4^+^ regulatory T cells, whose number was far more than that of B cells (Fig. S4D-S4F). In B6-Ighj KO mice, T cells were normal (Fig. S4C) and the infiltration of T lymphocytes after the first round of high-dose AAV injection was comparable to that in the wild-type mice (Fig. S4G-H), yet the redose of the 2^nd^ round of AAV showed significant GFP expression (Fig. 4N-O). Taken together, these results suggest that it is the neutralizing antibodies secreted by B cells, rather than the infiltration of T lymphocytes, that inhibit the redose of the same-serotype AAV in the brain.

### Lymphocytes infiltrate into the brain parenchyma through the perivascular space and the ventricle

After demonstrating that the high dose of AAV5 induced B cell infiltration and NAb production, we wondered how they infiltrated into the brain parenchyma. We first checked the integrity of the striatal BBB after high-dose AAV5 injection at different time points (7 dpi, 14 dpi, and 21 dpi). The structure of vasculatures (indicated by CD31 expression pattern and intensity, Fig S5A-B), the tight junction proteins (indicated by ZO-1, Fig. S5C-G) of the endothelial cells, the density of pericytes (indicated by PDGFRB, Fig. S5E-F), and glial endfeet (indicated by AQP4, Fig. S5G-K) were comparable to those in PBS-treated mice, suggesting that the BBB structure was relatively intact after high-dose AAV5 injection. Consistent with the structural intactness of BBB, the striatum receiving high dose of AAV5 injection exhibited little extravasation of FITC-d20 (molecular size, 20 kDa), which was comparable to that receiving PBS or without injection (Fig. S6L-S6N). Together, these analyses suggest that parenchymal administration of high-dose AAV5 did not break the integrity of BBB.

To dissect out the route of the infiltrated monocytes including B cells, we examined the distribution of the infiltrated cells in the brain parenchyma. Interestingly, the spreading pattern of CD45^+^ cells was quite unique, mostly located adjacent to the blood vessels, especially the large vessels (Fig. 5A). Among the blood vessels, we found that many CD45^+^ cells accumulated in the large blood vessels with diameters over 8 μm, while the number of CD45^+^ cells located near the small capillaries was quite limited (Fig. 5B). There is a perivascular space (PVS) between the blood vessels and glial limitans, which is easy to find in the postcapillary veins, serving as specialized immune niches in CNS where resides immune cells and collects the metabolite waste from interstital fluid from parenchyma.^11, 32^ When using GFAP to indicate the glial limitans along the blood vessels, we observed that many CD45^+^ cells as well as B220^+^ B cells located in the PVS between the astrocytic limitans and the blood vessels in the striatum following high dose AAV5 injection (Fig. 5C). Under normal condition, the number of immune cells within the PVS is quite few, but after damage, aging or CNS autoimmune disease (e.g. multiple sclerosis) immune cells increase dramatically through the comprmised BBB or CSF.^11, 32^ Since the BBB was not compromised by the high-dose AAV5, we checked the possibility of CSF as the source of the immune cells. Through the serial sections, we observed brain-wide distribution of the CD45^+^ cells along the rostral-caudal axis of ventricles (Fig. 5D). Enlarged images illustrated that both CD45^+^ cells and B220^+^ cells were dramatically elevated in the subventricular zone (SVZ) of lateral ventricle (LV, adjacent to the stritum where AAV injected) and the third ventricle (Fig. 5E-5F). Normally, ependymal cells in the ventricle form the brain-CSF barrier (BCSFB) to limit the invasion of immune cells from CSF to the brain parenchyma due to the tight-junctions.^11, 12, 13^ However, the intrastriatal injection of high-dose AAV5 triggered the CD45 positive cells to invade mainly along the ventricle. This phenomenon made us to suspect whether the BCSFB along the ventricle was compromised by the intrastriatal injection of high-dose AAV5. To explore the possiblity of infiltration through ventricles, we injected the fluorescent tracer OA647 (molecular weight: 45kDa) unilaterally into the right lateral ventricle 21 days after injecting high-dose AAV5 and analyzed the distribution of OA647 (Fig. 5G). 35 minutes after the OA647 injection is considered as a time point to study the influx of CSF.^38^ Under normal condition, the unlaterally injected tracer disperses to both sides of ventricle and flows into the glymphatic vessels with little leakage to the parenchymal tissue, as shown in the mouse brains receiving PBS (Fig. 5H, top row). However, in the brains that received the high-dose intrastriatal AAV5 injection, an increased signal of OA647 was observed in the parenchymal tissue around the ventricle (Fig. 5H, bottom row, and quantified in 5I), especially on the side receiving the injection of OA647 (Fig. S7D). Interestingly, the pattern of the leakage of the tracer was consistent with that of the infiltration of CD45^+^ cells (Fig. 5D and Fig. 5H). Additionally, the CXCL10 signal was enhanced in the astrocytes along the LV (Fig. S6E-S6F), further provide the chemotaxis for immune cells invasion through the LV. In contrast to the influx, the efflux of the CSF indicated by the distribution of OA647 examined at 120-minute after the intraventricular injection was comparable between the brains receiving PBS and high-dose AAV5 (Fig. S6A-C). Overall, these analyses indicated that high-dose AAV might increase the permeability of BCSFB along the ventricle nearby the injection sites and induce lymphocytes to invade the parenchyma through the perivascular space and ventricles.

**Figure 5.**
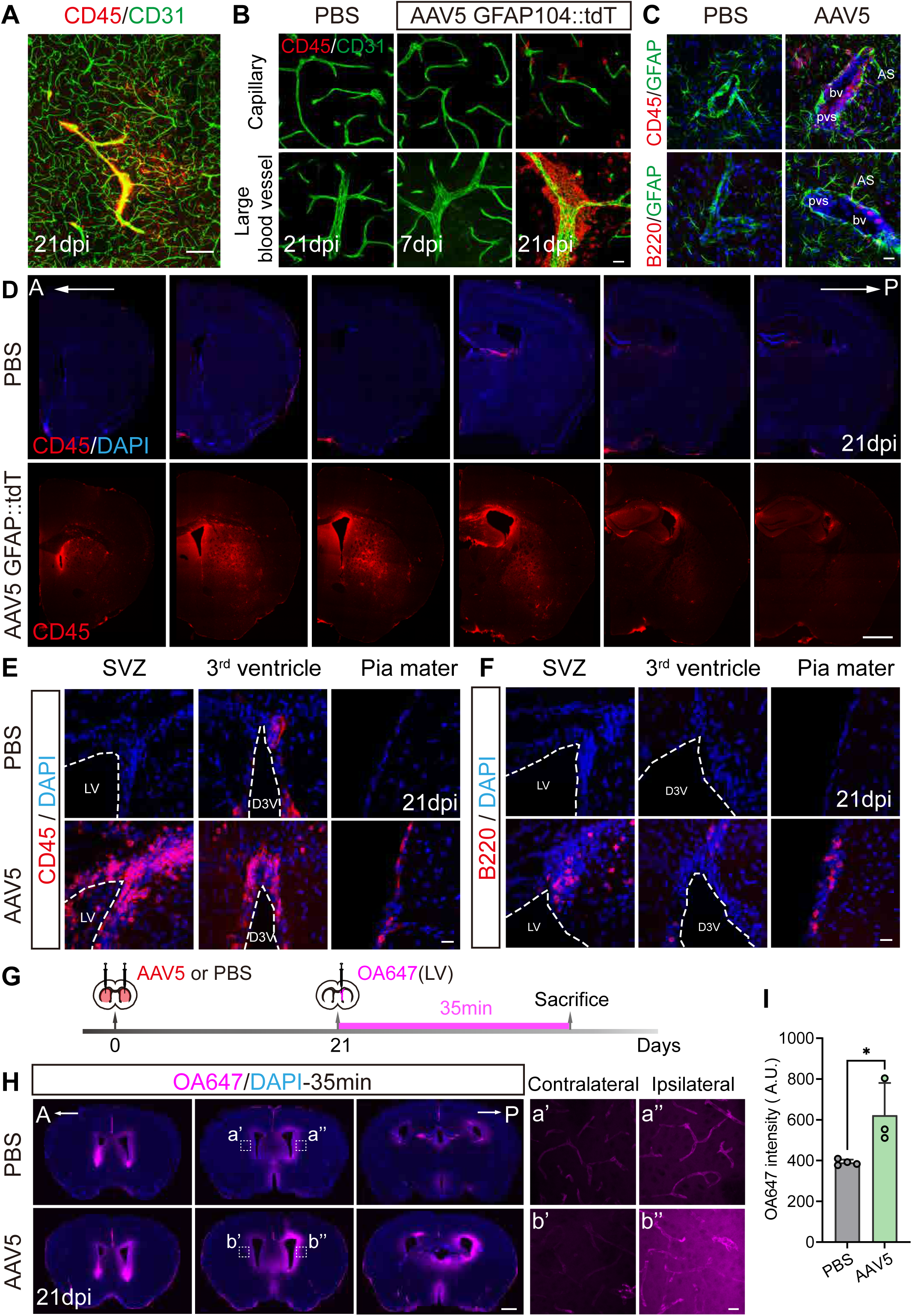
Lymphocytes infiltrate into the brain parenchyma through the perivascular space and the ventricles. (A) Low-magnification images showing that the location of the infiltrated lymphocytes (CD45^+^ cells in red) is associated with the blood vessels (CD31^+^ signal in green). Note that abundant CD45^+^ cells (red) are surrounding the blood vessels (indicated by CD31, green). Scale bar: 200 μm; dpi: days post injection. (B) High-magnification images displaying that the distribution of CD45^+^ cells in the striatum at different time points after receiving the intrastriatal injection of high-dose AAV5. Note that there is a robust invasion of CD45^+^ cells at 21 dpi, and majority of CD45^+^ cells accumulate around the large blood vessels (bottom panel), while a few CD45^+^ cells are located near the capillary (top panel). Scale bar: 20 μm; dpi: days post injection. (C) Representative images showing that the CD45^+^ cells (red, top panel) and B220^+^ cells (red, bottom panel) locate in the perivascular space outside the glial limitans (indicated by GFAP, green). The cell nuclei are counterstained with DAPI (blue). As, astrocyte; bv, blood vessels; PVS, perivascular space; scale bar: 20 μm. (D) Serial sections in the coronal plane revealing the brain-wide distribution of CD45^+^ cells (red) along the anterior-posterior axis at 21 days after intrastriatal injection of high-dose AAV5. The cell nuclei are counterstained with DAPI (blue). A: anterior; P: posterior; dpi: days post infection; dpi: days post infection; scale bar: 500 μm. (E-F) Representative images showing that both CD45^+^ (red, E) and B220^+^ (red, F) cells abundantly accumulate in the subventricular zone (SVZ) along the lateral ventricle, dorsal third ventricle (D3V), and pia mater. Ventricles are delineated by the white dashed lines. The cell nuclei are counterstained with DAPI (blue). Scale bars: 20 μm; dpi: days post injection. (G) Experimental design to assess ventricular system permeability with the tracer OA647. 21 days after PBS/AAV injection, OA647 (45 kDa) was injected into right lateral ventricle to circulate for 35 min. Then the mice were sacrificed for immunohistochemistry. (H) Images of serial sections in the coronal plane showing that 35 minutes after OA647 (magenta) injection, only little tracer leaks into the brain parenchyma that received PBS (top panel), while the signal OA647 significantly increases in the parenchyma around the ventricles (bottom panel) in the mouse that received high-dose intrastriatal AAV5 injection, especially on the ipsilateral side receiving the dye. a’, a’’, b’, and b’’ are high-magnification images of areas delineated by the white dashed squares in the low magnification images. The cell nuclei are counterstained with DAPI (blue). Scale bars: 1000 μm and 20 μm for a’, a’’, b’, and b’’; A: anterior; P: posterior; dpi: days post infection. (I) Quantitative data showing the OA647 intensity on the ipsilateral side where OA647 was injected is higher in the AAV5 group than that of the PBS group 35 minutes after dye injection. Mean (SD); two-way ANOVA with Sidak’s multiple comparison test, *P= 0.0296, n= 3 or 4 mice/group.

### Strategies to improve the re-administration of AAV after the high-dose injection

After illuminating the invasion of B cells in the brain and the production of NAbs induced by the high-dose AAV as a major barrier to the AAV re-administration, we searched for strategies to circumvent B cell responses to allow for redosing of AAV vectors. Firstly, we tried the approaches to diminish the effect of the neutralizing antibodies via injecting AAV5 empty capsids as decoys to capture antibodies through three different routes (retroorbital sinus, lateral ventricle, and striatum, Fig. S7A),^6^ but found that it failed to increase the second round viral infection (indicated by GFP signal, Fig. S7B and S7G). Then, we tried the intraventricular injection of IgG-degrading enzyme (IdeZ) that can rapidly and transiently cleave antibodies,^39^ but the expression of the second round of viral vectors (indicated by GFP signal) was still not improved (Fig. S7C, S7D, and S7H). When the strategies aiming to eliminate the parenchymal NAbs failed, we considered the approach to reduce the chemotaxis of B cell invasion. The inhibition of HDAC3 has been reported to reduce the expression of CXCL10 of the reactive astrocytes.^18^ Thus, HDAC3 inhibitor (RGFP966, 10 mg/kg) was daily administrated through intraperitoneal injection of 5 days after the first round of viral injection, and continued to the 3 days after the second round of virus injection (Fig. S7E). Unforturnately, the treatment of HDAC3 inhibitor did not increase the expression area of transgene (indicated by GFP signal) of the second round viral vectors (Fig. S7F and S7I). An alternative approach to circumvent antibodies induced by the first round of high-dose AAV was to switch the serotypes in the second round. We therefore tested high-dose AAV5 in the first round and then AAV9 in the second round (Fig. 6A), or high-dose AAV9 in the first round and then AAV5 in the second round (Fig. 6C). In both cases, the transgene expression of the 2^nd^ round AAV was not impaired (Fig. 6B, 6D, quantified in 6E-6F). This data suggests that changing the AAV serotypes enables effective re-administration of AAV. However, this strategy is costly for clinical applications as it would require GMP production and release testing of two different AAV products. As a matter of fact, another effective but simple approach to enable the AAV redose was to reduce the dose of AAV in the first round of intracraninal injection (Fig. 1A, 1C, 1H and 1I). Together, the above results implicated that it might not be easy to eliminate the NAbs once the B cells infiltrated into the CNS parenchymal tissues. It is important to find a minimal effective dosage in the first round of AAV gene therapy to minimize the B cell invasion and NAb production if redosing is a desirable option.

**Figure 6.**
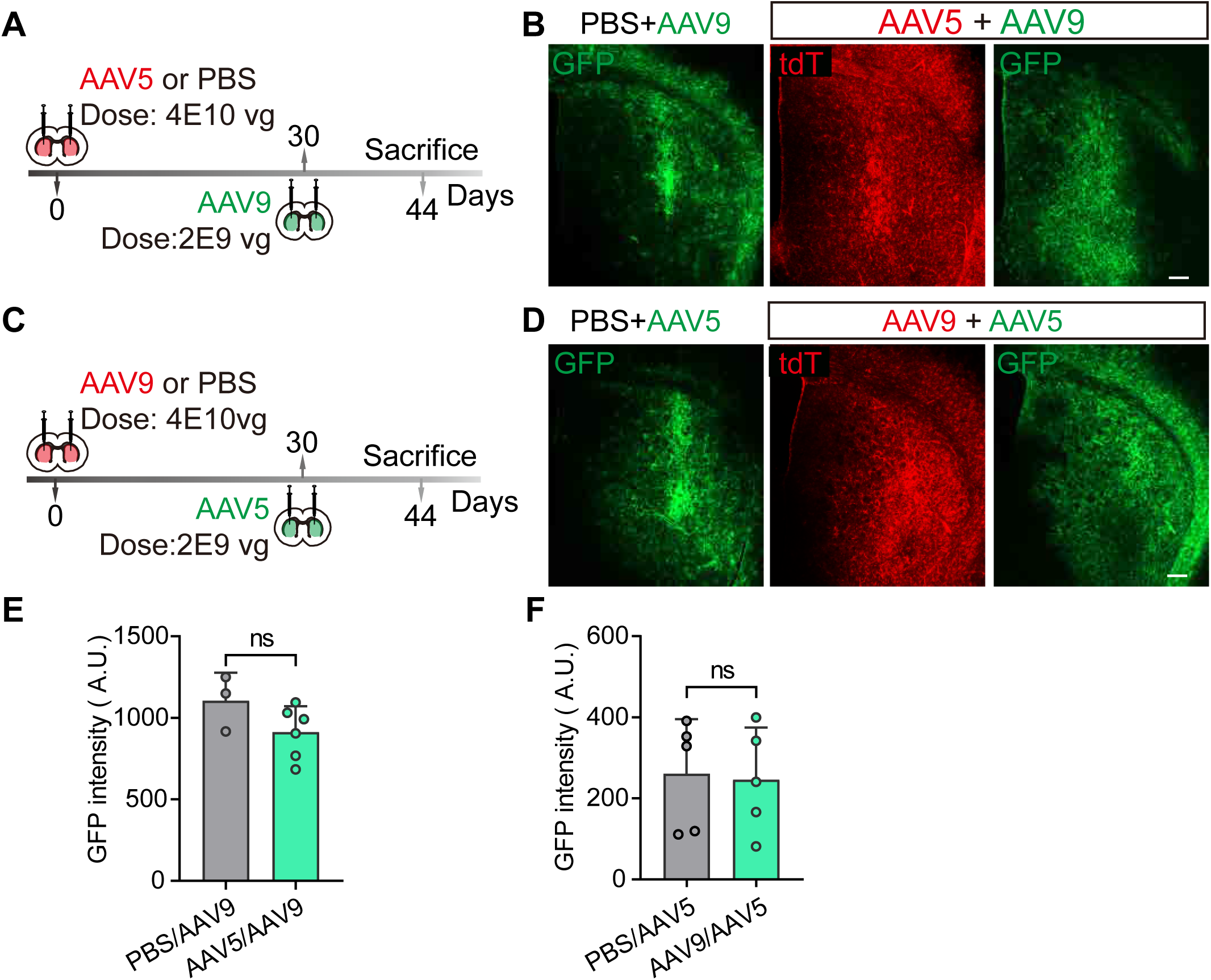
Strategies to improve the re-administration of AAV after the high-dose injection. (A) Experimental design to use different AAV serotypes to circumvent NAbs. AAV9 GFAP104::GFP was injected into the same site 30 days after the intrastriatal administration of AAV5 GFAP104::tdTomato. GFP and tdTomato expression were examined 2 weeks after the second injection. Mice receving intrastriatal injection of PBS in the first round were used as a control. (B) Typical images showing robust GFP expression in the striatum, suggesting that the expression of transgene delivered by AAV9 (GFP, green) in the second round is not affected by high dosage of AAV5 (tdT, red). Scale bar: 200 μm (C) Experimental design to reverse the injection order of the serotypes of the two rounds: AAV9 expressing tdTomato was used as the first round and AAV5 expressing GFP was used as the second round. (D) Representative images showing that the expression of the transgene (GFP, green) delivered by AAV5 in the second round is not inhibited by high-dosage AAV9 (tdT, red) in the first round. Scale bars: 200 μm (E-F) Quantitation showing that the expression of the second-round AAV (GFP) in the striatum that received the high-dose AAV in the first round is comparable to that received PBS in the first round if the AAV serotypes used in the two rounds are different, no matter if the first round is AAV5 (H) or AAV9 (I). Mean (SD); unpaired t test, n= 3 or 5 mice/group.

## Discussion

Neutralizing antibodies (NAbs) is considered as the major obstacle to the re-administration of rAAV when targeting the peripheral tissues,^6, 40^ but whether it impedes the rAAV redosing in the central nervous system has not been fully elucidated . In this study, we show that intrastriatal injection of rAAV induces the invasion of B lymphocytes into the parenchymal tissue and the production of NAbs in a dose-dependent manner, which compromises the re-administration of rAAV of the same serotype. Such immune-suppression effect of high-dose rAAV lasts at least 5 months and spreads widely in brain regions beyond the injection site. The high-dose intracranial injection of rAAV5 does not disrupt the integrity of BBB, but rather increases the permeability of brain-CSF barrier (BCSFB) along the ventricles. In addition, it activates the local glial cells in the injection regions to evelate the expression of chemokines CXCL10 and CXCL9, and recruit the immune cells including B cells to infiltrate into the parenchyma through the specialized immunological compartments in the brain including perivascular space and ventricles. Change of the serotype of the rAAV of the second injection or genetic depletion of the B lymphocytes can circumvent the immunosuppression of the high-dose rAAV. This is the first study showing a dosage dependent activation of adaptive immune in the adult mouse brain after intracranial rAAV injection, and a critical role of B lymphocyte infiltration through the border specialized immune niches of CNS.

Previous studies on the compromising effect of intracranial injection of rAAVs after redosing are controversial. Peden et al. reported that the striatal administration of rAAV2 induced an immune response against the following redosing. But this is a short-term immunosuppression within 11 weeks.^9^ On the contrary, Wang et al. found that a thalamic injection of rAAV9 did not change the transduction capability of the following intracranial injection of rAAV9 with a 5-week interval.^10^ On the other hand, pre-immunization with rAAV9 by intramuscular injection significantly reduced the transduction of rAAV9 in the brain regions.^10^ Despite this controversy, both studies agreed that intracranial injection of rAAVs did not induce the production of NAbs and the compromising effect on redosing might be pleiotropic immune response beyond generating circulating NAbs.^9, 10^ Here, our study unambiguously demonstrates that the dosage dependent compromising effect of the intracranial injection of rAAVs on the redosing of rAAV is due to the production of NAbs by infiltrated B cells. The NAbs is present in the parenchymal tissues in both injection regions and non-injection regions, and the titer increases over time after the injection. The time course of the parenchymal NAbs is consistent with the time course of the compromising effect. Interestingly, the high-dose intracranial injection of rAAVs induces the NAb generation both in the sera and in the brain tissues, but the generation of serum NAbs is faster than that of parenchymal NAbs. This phenomenon is consistent with the previous study reporting a delayed production of NAbs in the mouse CNS after the infection of the neurotropic JHM strain of mouse hepatitis virus (JHMV).^41^ In that study, virus-specific Ab-secreting cells (ASC) peak within the CNS by day 21 post injection.^41^ In this study, we also discovered ASC cell infiltration in the injection side at 21 dpi. The compromising effects of high-dose rAAV lasted at least 5 months, which is contrast to the previous report of short-term effect within 11 weeks by Peden et al.,^9^ but consistent with the observation of long-term presence of virus-induced ASCs within CNS at least 90 days after JHMV infection.^41^ Our study implies that the activation of adaptive immune response induced by the intracranial administration of the engineered rAAVs may be long-term and may not fade away easily.

When we investigated the ASC infiltration and the production of NAbs, we firstly considered the possibility of BBB breakdown . A recent study reported the disruption of BBB resulting from the high-titer rAAV intracranial injection, leading to the GZMB^+^ T lymphocyte infiltration and neuronal damage.^42^ In our study, we also observed the massive lymphocytes including ASC invasion in the injection sites after the intrastriatal injection of high-dose rAAV. We initially thought that the intergrity breakdown of BBB could be an easy explanation of the presence NAbs in both the injection sites and non-injection sites. However, bio-informatic analysis based on RNA-seq data provided little clues on the BBB breakdown (e.g. upregulation of *Icam1*, *Mmp3* and *Mmp13* reported previously) after the intrastriatal administration of high-dose AAV5.^42^ In addition, both the histological analyses and the function assays through the intravenous injection of fluorescent dye did not support the damage of the BBB. The discrepancy in the disruption of BBB between the previous and this study may be attributed to complicated reasons intertwined with the selection of promoters, the serotypes and the injection sites. The previous study observed the BBB breakdown in the cortex when they used a long version of *GFAP* promoter (*Gfa2*, with a length of 2.2 kb) with AAV9 serotype,^42^ whereas we used a short version of Gfa2 (GfaABC1D, with a length of 681) promoter and AAV5 serotype to target astrocytes in the striatum. Although there was the infiltration of T lymphocytes expressing GZMB in our study, the density of local neurons in the striatum was normal after the intracranial microinjection of high-dose rAAV5 (data not shown). Consistently, the total genes changed after the intrastriatal injections of high-dose rAAV5 GFAP104::GFP in this study is only 1/7^th^ of that after intracerebral injection of high titer rAAV9 GFAP2.2::GFP in previous study.^42^ Besides the breakdown of BBB, intracranial administration of high-dose rAAVs has also been reported to disrupt the neural circuits.^43, 44^ In one study using the same serotype (AAV5) and similar viral vector (GFAP104::GFP) to this study, they found that microinjection of high-dose rAAV5 in the hippocampus selectively induce astrocytic gliosis that downregulated astrocytic expression of glutamine synthetase (GS) to cause the deficits in neuronal inhibition.^43^ In this study, the intrastriatal delivery of high-dose rAAV5 also activated the astrocytes, but the GS expression was not altered (data not shown). Together, all these results indicate that the adverse effects induced by intracranial injection of high-dose rAAVs depend on the viral vector, the serotype and the injection regions.

If the BBB is intact, how do the B lymphocytes infiltrate into the CNS to produce NAbs? Recent studies have identified special niches in the brain’s border, including the choroid plexus (CP), meninges, and the perivascular space, where various innate and adaptive immune cells populate in the surveillance and defense of the brain.^11, 12^ The blood vessels in meninges and CP are fenestrated, allowing the circulating immune cells to migrate inwards to CSF.^11, 12, 32^ In addition, the skull bone marrow provides rapid access of immune cells to the brain through microscopic channels within the skull,^45, 46^ and the brain-draining cervical lymph nodes (CLNs) collect lymphatic fluid from the brain through the meningeal lymphatic system.^47, 48, 49, 50, 51, 52^

Both of them can sense the CNS antigens and motivate the immune cells to enter the CNS through the specialized immune niches in the border.^11, 12^ In this study, we noticed that most of the immune cells (including T lymphocytes, B lymphocytes, and ASCs) accumulated around the perivascular spaces (PVS) and along the ventricle regions adjacent to the injection site, suggesting that they potentially migrate from these specialized niches at the border of CNS. The permeability of the brain-CSF barrier (BCSFB) along the ventricular regions adjacent to the injection sites is increased after the intracranial injection of high-dose rAAVs, which suggests a compromise of BCSFB in this region and provides a route for the invasion of ASCs within the CSF. The compromise of BCSFB along the ventricle by high-dose rAAVs has not been reported before, expanding our understanding of the adverse effects of high-dose rAAVs to the CNS. In addition to PVS and ventricles, the number of immune cells in the pial matter also increased after the high-dose rAAV injection. Pial matter is a component of lepitomeninge, which locates beneath the dura matter and where CSF flows,^13^ providing additional routes for the invasion of immune cells including B lymphocytes in the CNS.

Previous studies report the retention of ASCs within the CNS by increasing the CXCL9/10-CXCR3 axis for recruitment and BAFF/CXCL12 for survival in the autoimmune disease (MS) and neurotrophic viral infections in the CNS.^20, 24, 29, 36, 41, 53^ In this study, the augmentation of CXCL9, CXCL10 and *Cxcl12* was also observed in the striatum after the microinjection of high-dose rAAV. The CXCL10 is expressed exclusively in the reactivated astrocytes adjacent to the PVS and lateral ventricle where the B lymphocytes accummulate. The signal of CXCL9 is found in astrocytes and cells associated with the PVS. CXCL9, CXCL10 and CXCL12 are chemockines to recruitment T lymphocytes.^19, 54^ CD4^+^ regulatory T lymphocytes regulate the B lymphocyte maturation and formation of the memory B cells.^19, 54^ Although we prove in this study that the infiltration of T lymphocytes is not the direct executor to compromise the AAV redosing, whether they facilitate the formation of memory B cells requires further investigation.

This study also has some limitations. One is that although we demonstrate the infiltration of ASCs and the production of NAbs as the major hurdle of rAAV redosing, we have not been able to develop a B cell-focused immune suppression protocol for efficient AAV readministration to the brain. OCREVUS^®^ (ocrelizumab) is a FDA-approved therapeutic monoclonal antibody against CD20-positive B cells to treat relapsing or primary progressive forms of multiple sclerosis (MS).^55, 56, 57^ Recent study developed a protocol of using a combination of monoclonal Ab therapy against CD20 (for B cell depletion) and BAFF (to slow B cell repopulation) to prevent immune responses and abrogate prolonged Ab formation, which elevated the redosing efficiency in hepatic rAAV transfer.^40^ Thus, the utilization of monoclonal Abs against B cells might be a potential strategy to increase the efficiency of intracranial redosing of rAAVs. However, considering the relative integrity of BBB after intracranial injection of high-dose rAAV, whether the monoclonal Abs can effectively penetrate the brain tissue requires further investigation. In this study, employing empty capsid decoys or IgG-degrading enzyme, which worked efficiently in the peripheral system,^6,39^ failed to attenuate the compromising effect of the high-dose rAAVs in the brain.

Another limitation is that we cannot rule out the possibility of a transient leakge of the BBB after the high-dose AAV infection. In this experiment we mainly checked 21 dpi when the infiltration of immune cells was evident. Whether the BBB is temporarily compromised before rehealing again requires further study.

In conclusion, for the first time, this study reveals that intrastriatal administration of high-dose rAAV activates local glial cells to elevate the chemokines and increases the permeability of brain-cerebrospinal fluid barrier (BCSFB), to recruit B lymphocytes from the PVS and ventricle regions and the production of NAbs. This induces adaptive immune responses that compromise the second-round intraparenchymal rAAV redosing of the same serotype. Although switching the serotype of rAAV in the second round of intracranianl injection enables the redosing, the more effective approach is to find a minimal effective dosage in the first round to minimize the B cell invasion in the CNS.

## Materials and Methods

### Animals

Adult C57BL/6J mice (> 8 weeks, Guangdong Yaokang Biotechnology, China) and B6-Ighj KO mice (C001344, https://www.cyagen.com/cn/zh-cn/business/drug-filter-evaluate-mouse/immunodeficiency-mouse/B6-Ighj-KO.html,> 8 weeks, Cyagen Guangzhou Biosciences, China) were used in this experiment. Both male and female mice were used in this study and housed under standard conditions (room temperature 22°C-25°C, humidity 40% - 60%, a 12-h light-dark cycle) with sufficient water and food. Experimental protocols were approved by the Laboratory Animal Ethics Committee of Jinan University (approval No. IACUC- 20230411-02).

### AAV vector production

The AAV5 and AAV9 used in this experiment were constructed and produced by PackGene Biotech (Guangzhou, China). All rAAVs were produced by co-transfection of a large number of ultrapure plasmids with HEK293 cells, and the rAAV particles were concentrated and purified by ultrahigh-speed gradient centrifugation. Genome titer and AAV purity were determined by SYBR Green qPCR and SDS-PAGE electrophoresis. The empty shell AAV5 virus was prepared in the same way, with the exceptation to exclude the genome plasmid of the vector (cis-acting plasmid) during the transfection process. The titer of the empty AAV9 viral capsids was determined using an AAV9 Titration ELISA kit (PRAAV9, Progen). AAV titers were diluted with PBS containing 0.001% F-68 before use. All AAVs were stored in an 80°C freezer and used within 12 months of production. Details of the AAVs used in this study are shown as follows:

| Virus | Experimental titer |
| --- | --- |
| AAV5 GFAP104::tdTomato | 1E3 vg/mL or 5E11 vg/mL |
| AAV5 GFAP104::GFP | 1E3 vg/mL or 5E11 vg/mL |
| AAV5 Syn::mCherry | 1E3 vg/mL |
| AAV5 Syn::GFP | 5E11 vg/mL |
| AAV5 CAG::Luciferase | 5E9 vg/mL or 5E8 vg/mL |
| AAV5 empty capsid | 1E3 vg/mL |
| AAV9 Syn::mCherry | 1E3 vg/mL |
| AAV9 Syn::GFP | 5E11 vg/mL |
| AAV9 GFAP104::tdTomato | 1E3 vg/mL or 5E11 vg/mL |
| AAV9 GFAP104::GFP | 1E3 vg/mL or 5E11 vg/mL |

### Stereotaxic viral injection

All mice were anesthetized by intraperitoneal injection of Avertin (20 mg/kg of 1.25% solution, Sigma, T48402). After shaving and disinfection, artificial eye ointment was applied to protect the eyes of the mice. The scalp was incised along the midline of the mouse brain, and a hole (0.5 mm) was drilled above the skull. With the bregma as the origin, the rAAV preparation was then delivered using glass microelectrode at 200 nL/min into the striatum (AP: + 1.0 mm, ML: ± 2.0 mm, DV: - 3.0 mm, 2µL), hippocampus (AP: - 2.1 mm, ML: ± 1.8 mm, DV: - 1.5 mm, 1.5 µL), cerebellum (AP: - 6.1 mm, ML: ± 1.5 mm, DV: - 2.0 mm, - 2.8 mm, 1 µL),or lateral ventricle (AP: -0.3 mm, ML: ± 0.9 mm, DV: - 2.0 mm, 5 µL). After virus injection, the glass microelectrode was left in the parenchyma for about 10 minutes and then slowly withdrawn.

### Tissue preparation and immunofluorescence

The anesthetized mice were trans-cardially perfused with ice-cold saline solution. The brains were quickly dissected and fixed in 4% paraformaldehyde (PFA) overnight and then cut into 30 µm sections using a vibrating microtome (Leica, VTS1200, Germany). For immunostaining, the brain slices were firstly washed 3 times in phosphate buffered saline (PBS, pH: 7.35) and then incubated in blocking solution (5% donkey serum + 0.3% TritonX-100) for 1 h at room temperature. Primary antibodies were dissolved in blocking solution (Table S1: Primary antibody information) and then incubated with primary antibodies overnight at 4°C. Next, the brain slices were washed 3 times with PBS for 15 minutes each. Afterwards, the samples were incubated with fluorescently labeled secondary antibodies (Alexa Flour 448, 1:1000; Alexa Flour 555,1:1000; Alexa Flour 647, 1:500; Life technologies) and DAPI (1:2000) for 2 hours and washed with extensive PBS at room temperature. Later, the sections were mounted on glass slide with anti-fade mounting medium (Sigma-Aldrich, 10981).

### qRT-PCR

The mice were trans-cardially perfused with ice-cold saline and then quickly dissected the striatum and hippocampus into 2 mL EP tubes. Tissue was homogenized in DNA/RNA Lysis Buffer using an ultrasonic processor (Vibra-Cell™, Sonics, U.S.A). DNA and RNA were then extracted according to Quick-DNA/RNA™ Miniprep Kit (Cat. #D7001). cDNA synthesis was performed using PrimeScript™ RT reagent Kit (Cat. #RR037A). After collecting DNA and cDNA, qRT-PCR was performed with SYBR green fluorescent PCR kit (Cat. # 208054) and ran on a CFX96 RealTime System (Bio-Rad Laboratories). The relative expression level was calculated as 2(-ΔΔCT). The primers used for PCR are listed as follows:

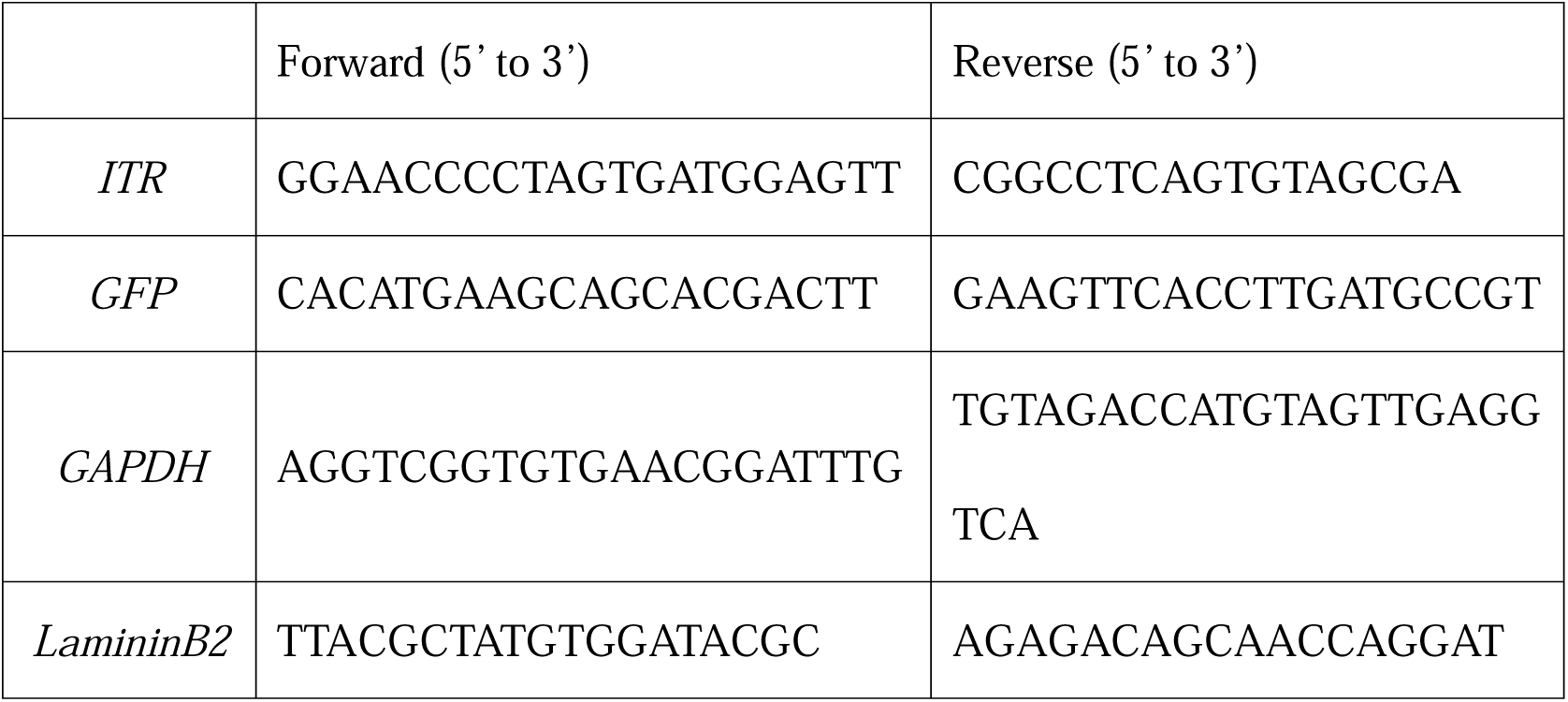

### *In Vitro* AAV- NAb Assay Tissue collection

Mice were trans-cardially perfused with ice-cold saline solution. The brains were quickly harvested to carefully dissect the striatum and hippocampus on ice bricks. Tissue were stored at 80°C until Nab assay.

### AAV-NAb Assay

Primary rat astrocytes were dissociated and cultured as described previously.^58^ 24 hours before the *in vitro* NAb assay, the cells were seeded on well of 96-well plate at the density of 2,500 cells/well. The frozen tissue was minced and sonicated in RAPI lysis buffer (1 mg/10 µL, Cat. #P10013B) with PMSF protease inhibitor (Beyotime, Cat #ST506). The suspension was then centrifuged (12000 rpm, 4°C) for 30 minutes and the supernatant was collected. The total protein concentration in the clarified lysate was then quantified using the BCA assay (Cat. #PC0020). Serum and tissue samples were then heat-inactivated at 56°C for 30 minutes, and then serially diluted 4-fold using Dulbecco’s Modified Eagle’s Medium (DMEM) as a diluent. AAV5 CAG::Luciferase was added into the dilutions containing the serum sample (5E9GC/mL) and tissue sample (5E8GC/mL). After incubated at 37°C/5% CO_2_ incubator (Thermofisher) for 30 minutes later, the pre-incubated virus/tissue mixtures were added to each well containing rat astrocytes (2500 cells/well) and cultured overnight at 37 °C /5% CO_2_. The virus-containing medium was then removed and replaced with normal glial medium. The medium was then removed every 48 hours. Triplicate wells were used for each dilution of the sample. Five days after addition of the virus, the culture medium was washed off with PBS, astrocyte cultures were lysed with tissue lysis buffer, and Luc in the tissue lysate was quantified using the Lysol Luciferase Assay System (Promega, Cat. #E1501). The transduction inhibitory effect of each dilution of the test sample was calculated by comparison with astrocyte lysates incubated in normal glial medium. The range between the lowest dilution factor that yielded more than 50% transduction inhibition and the highest dilution factor that could not inhibit transduction by more than 50% was defined as the NAb titer.

### Bulk RNA-seq and analysis

Mice were perfused with ice-cold saline solution. Next, the striatum and hippocampus were quickly collected into 2 mL pre-cooled EP tubes and frozen in liquid nitrogen for 30 min, and then stored at 80°C for RNA extraction and bioinformatics analysis. RNA extraction was performed by Gene Denovo Biotechnology Co., Ltd (Guangzhou, China), and RNA purity and concentration were detected by NanoPhotometer spectrophotometer and Qubit2.0 Fluorometer. RNA library sequencing was performed on the Illumina HiseqTM 2500/4000 by Gene Denovo Biotechnology. Bioinformatic analysis was performed on the real-time interactive online data analysis platform Omicsmart (http://www.omicsmart.com). Genes/transcripts with FDR < 0.05 and difference fold change ≥ 1.5 were considered as differentially expressed genes/transcripts.

### Intraventricular tracer injection

Ovalbumin Alexa Fluor 647 (OA647, Invitrogen) was constituted in artificial CSF at a concentration of 0.5%. For intraventricular tracer injection, OA647 was injected into the right lateral ventricle (200 nL/min, 6 µL) with a 30 GA steel needle. 35 minutes or 120 minutes later, the animal were perfused with 4% ice-cold PFA and the brain were harvested. The brains were post-fixed overnight and then cut into 60 µm sections on a vibratome.

### Blood-brain barrier permeablity assay

After the injection of AAV5 or PBS into the striatum for 21 days, all mice received 100 μL 20-kDa dextran-FITC (20 mg/mL) via the retroorbital venous sinus. 2 hours later, the mice were perfused with 4% ice PFA. The brains and livers were rapidly collected, and incubated in 4% PFA overnight at 4 °C. Brains were cut into 60 µm sections using a vibrating microtome. The livers were dehydrated with 20%, 30%, and 30% sucrose solution until the tissue sank. After the embedment in Optimal Cutting Temperature (Tissue-Tek® O.C.T.), livers were cut into 50 µm sections using a crystat (Cryostar NX50, Thermo scientific, Germany).

### Drugs to improve AAV re-administration

RGFP966 (10 mg/kg, Repligen, Cat. #S7229) was dissolved in DMSO and diluted in a vehicle of 40% (vol/vol) PEG300 (Selleckchem, Cat. #S6704), 10% Tween80 and 45% ddH2O, the final DMSO concentration was 5% (vol/vol) for the vehicle. RGFP966 was injected intraperitoneally once a day.

IdeZ (2700U/kg, New England Biolabs, Cat. #P0770S) was dissolved in saline and injected into the bilateral lateral ventricles (10µL) with a glass microelectrode.

### Statistical analysis

All images were captured with a Carl Zeiss microscope (Imager Z2, Carl Zeiss, Germany), a SlideView VS200 (VS200, Olympus, Japan), or a Zeiss confocal microscope (LSM 880; Carl Zeiss, Germany). For quantitative analysis, three brain sections per animal were selected for counting. For quantification of fluorescence intensity and area, all counting was performed by Zeiss ZEN 2.3 (Blue Edition, Göttingen, Germany) or ImageJ, and the background intensity was subtracted from the fluorescence intensity. For cell density quantification, 5 or 8 areas per brain section were randomly selected for quantification (×20 lens for 5 regions, ×40 lens for 8 regions). For calculating perivascular AQP4 polarity, fluorescence intensity values were recorded at 80 different pixel locations across 10 penetrating blood vessels in the mouse striatum, and the quantification method was adopted from a previous study (ref). Data are shown as mean (SD). Statistical analysis was performed using unpaired Student’s t-test (two-tailed Mann-Whitney test) for two-group comparison, one-way ANOVA (Sidak’s multiple comparison test) for multiple group comparisons, and two-way ANOVA analysis (Sidak’s multiple comparison test) for multivariate and multiple group comparisons. P value over 0.05 is considered as statistical significance.

## Data availability statement

The data that support the findings of this study are available from the corresponding author upon reasonable request.

## Acknoledgments

This study is supported by Science and Technology Projects of Guangzhou, No. 202206060002 (to G.C. and W.L); Natural Science Foundation of Guangdong Province of China, No. 2023A1515011719 (to W.L.); the Pearl River Innovation and Entrepreneurship Team, No. 2021ZT09Y552 (to G.C.).

## Author Contributions

W.L., Y-G.X, and G.C. conceptulized the research. Y-G.X., X-N.B. performed all experiments with the assistance of J-H.L., S-H.W., Y-Y.W, and H-L.L.. Y-G.X. and S-G.L. performed bioinformatics analysis. Y-G.X., X-N.B., and K.L. performed neutralizing antibody detection and qPCR. Y-G.X. and W.L. analyzed the data and made the figures. W.L., Y-G.X. and G.C. wrote the manuscript and all authors discussed during and after the writing.

## Reference

1. Li C, Samulski RJ. Engineering adeno-associated virus vectors for gene therapy. Nature reviews 21, 255–272 (2020).

2. Gao J, et al. Gene therapy for CNS disorders: modalities, delivery and translational challenges. Nature Reviews Neuroscience, (2024).

3. Wang JH, Gessler DJ, Zhan W, Gallagher TL, Gao G. Adeno-associated virus as a delivery vector for gene therapy of human diseases. Signal Transduct Target Ther 9, 78 (2024).

4. Chu WS, Ng J. Immunomodulation in Administration of rAAV: Preclinical and Clinical Adjuvant Pharmacotherapies. Front Immunol 12, 658038 (2021).

5. Muhuri M, et al. Overcoming innate immune barriers that impede AAV gene therapy vectors. J Clin Invest 131, (2021).

6. Ertl HCJ. Circumventing B Cell Responses to Allow for Redosing of Adeno-Associated Virus Vectors. Human gene therapy, (2023).

7. Mendell JR, et al. Testing preexisting antibodies prior to AAV gene transfer therapy: rationale, lessons and future considerations. Molecular Therapy - Methods & Clinical Development 25, 74–83 (2022).

8. Mastakov MY, Baer K, Symes CW, Leichtlein CB, Kotin RM, During MJ. Immunological aspects of recombinant adeno-associated virus delivery to the mammalian brain. Journal of virology 76, 8446–8454 (2002).

9. Peden CS, et al. Striatal readministration of rAAV vectors reveals an immune response against AAV2 capsids that can be circumvented. Mol Ther 17, 524–537 (2009).

10. Wang D, et al. Adeno-Associated Virus Neutralizing Antibodies in Large Animals and Their Impact on Brain Intraparenchymal Gene Transfer. Molecular Therapy - Methods & Clinical Development 11, 65–72 (2018).

11. Castellani G, Croese T, Peralta Ramos JM, Schwartz M. Transforming the understanding of brain immunity. Science 380, eabo7649 (2023).

12. Goertz JE, Garcia-Bonilla L, Iadecola C, Anrather J. Immune compartments at the brain’s borders in health and neurovascular diseases. Semin Immunopathol 45, 437–449 (2023).

13. Rua R, McGavern DB. Advances in Meningeal Immunity. Trends in molecular medicine 24, 542–559 (2018).

14. Lee Y, Messing A, Su M, Brenner M. GFAP promoter elements required for region-specific and astrocyte-specific expression. Glia 56, 481–493 (2008).

15. Perea G, Yang A, Boyden ES, Sur M. Optogenetic astrocyte activation modulates response selectivity of visual cortex neurons in vivo. Nature communications 5, 3262 (2014).

16. Unterholzner L, et al. IFI16 is an innate immune sensor for intracellular DNA. Nature immunology 11, 997–1004 (2010).

17. Ning S, Pagano JS, Barber GN. IRF7: activation, regulation, modification and function. Genes & Immunity 12, 399–414 (2011).

18. Clayton BLL, et al. A phenotypic screening platform for identifying chemical modulators of astrocyte reactivity. Nat Neurosci, (2024).

19. Lam N, Lee Y, Farber DL. A guide to adaptive immune memory. Nature Reviews Immunology, (2024).

20. Tokunaga R, et al. CXCL9, CXCL10, CXCL11/CXCR3 axis for immune activation - A target for novel cancer therapy. Cancer Treat Rev **63**, 40-47 (2018).

21. Greenhalgh AD, David S, Bennett FC. Immune cell regulation of glia during CNS injury and disease. Nature reviews Neuroscience 21, 139–152 (2020).

22. Sofroniew MV. Astrocyte Reactivity: Subtypes, States, and Functions in CNS Innate Immunity. Trends Immunol 41, 758–770 (2020).

23. Xu S, Lu J, Shao A, Zhang JH, Zhang J. Glial Cells: Role of the Immune Response in Ischemic Stroke. Front Immunol 11, 294 (2020).

24. Graver JC, et al. Association of the CXCL9-CXCR3 and CXCL13-CXCR5 axes with B-cell trafficking in giant cell arteritis and polymyalgia rheumatica. J Autoimmun 123, 102684 (2021).

25. Han RT, Kim RD, Molofsky AV, Liddelow SA. Astrocyte-immune cell interactions in physiology and pathology. Immunity 54, 211–224 (2021).

26. Niso-Santano M, Fuentes JM, Galluzzi L. Immunological aspects of central neurodegeneration. Cell Discov 10, 41 (2024).

27. Chistiakov DA, Killingsworth MC, Myasoedova VA, Orekhov AN, Bobryshev YV. CD68/macrosialin: not just a histochemical marker. Lab Invest 97, 4–13 (2017).

28. Linnerbauer M, Wheeler MA, Quintana FJ. Astrocyte Crosstalk in CNS Inflammation. Neuron 108, 608–622 (2020).

29. Phares TW, Stohlman SA, Hinton DR, Bergmann CC. Astrocyte-derived CXCL10 drives accumulation of antibody-secreting cells in the central nervous system during viral encephalomyelitis. Journal of virology 87, 3382–3392 (2013).

30. Rahmat-Zaie R, Amini J, Haddadi M, Beyer C, Sanadgol N, Zendedel A. TNF-α/STAT1/CXCL10 mutual inflammatory axis that contributes to the pathogenesis of experimental models of multiple sclerosis: A promising signaling pathway for targeted therapies. Cytokine 168, 156235 (2023).

31. Vanlandewijck M, et al. A molecular atlas of cell types and zonation in the brain vasculature. Nature 554, 475–480 (2018).

32. Mastorakos P, McGavern D. The anatomy and immunology of vasculature in the central nervous system. *Sci Immunol* 4, (2019).

33. Altin JG, Sloan EK. The role of CD45 and CD45-associated molecules in T cell activation. Immunol Cell Biol 75, 430–445 (1997).

34. Hermiston ML, Xu Z, Weiss A. CD45: A Critical Regulator of Signaling Thresholds in Immune Cells. Annual Review of Immunology 21, 107–137 (2003).

35. Cardani-Boulton A, Boylan BT, Stetsenko V, Bergmann CC. B cells going viral in the CNS: Dynamics, complexities, and functions of B cells responding to viral encephalitis. Immunol Rev 311, 75–89 (2022).

36. Rodriguez-Mogeda C, Rodríguez-Lorenzo S, Attia J, van Horssen J, Witte ME, de Vries HE. Breaching Brain Barriers: B Cell Migration in Multiple Sclerosis. Biomolecules 12, (2022).

37. Gu H, Zou YR, Rajewsky K. Independent control of immunoglobulin switch recombination at individual switch regions evidenced through Cre-loxP-mediated gene targeting. Cell 73, 1155–1164 (1993).

38. Iliff JJ, et al. A paravascular pathway facilitates CSF flow through the brain parenchyma and the clearance of interstitial solutes, including amyloid β. Sci Transl Med 4, 147ra111 (2012).

39. Elmore ZC, Oh DK, Simon KE, Fanous MM, Asokan A. Rescuing AAV gene transfer from neutralizing antibodies with an IgG-degrading enzyme. JCI Insight 5, (2020).

40. Rana J, et al. B cell focused transient immune suppression protocol for efficient AAV readministration to the liver. Molecular therapy Methods & clinical development 32, 101216 (2024).

41. Tschen SI, Stohlman SA, Ramakrishna C, Hinton DR, Atkinson RD, Bergmann CC. CNS viral infection diverts homing of antibody-secreting cells from lymphoid organs to the CNS. Eur J Immunol 36, 603–612 (2006).

42. Guo Y, et al. High-titer AAV disrupts cerebrovascular integrity and induces lymphocyte infiltration in adult mouse brain. Molecular therapy Methods & clinical development 31, 101102 (2023).

43. Ortinski PI, et al. Selective induction of astrocytic gliosis generates deficits in neuronal inhibition. Nat Neurosci 13, 584–591 (2010).

44. Suriano CM, et al. An innate immune response to adeno-associated virus genomes decreases cortical dendritic complexity and disrupts synaptic transmission. Mol Ther, (2024).

45. Cugurra A, et al. Skull and vertebral bone marrow are myeloid cell reservoirs for the meninges and CNS parenchyma. Science 373, (2021).

46. Mazzitelli JA, et al. Cerebrospinal fluid regulates skull bone marrow niches via direct access through dural channels. Nat Neurosci 25, 555–560 (2022).

47. Louveau A, et al. Structural and functional features of central nervous system lymphatic vessels. Nature 523, 337–341 (2015).

48. Louveau A, Da Mesquita S, Kipnis J. Lymphatics in Neurological Disorders: A Neuro-Lympho-Vascular Component of Multiple Sclerosis and Alzheimer’s Disease? Neuron 91, 957–973 (2016).

49. Ahn JH, et al. Meningeal lymphatic vessels at the skull base drain cerebrospinal fluid. Nature 572, 62–66 (2019).

50. Li X, et al. Meningeal lymphatic vessels mediate neurotropic viral drainage from the central nervous system. Nat Neurosci 25, 577–587 (2022).

51. Smyth LCD, et al. Identification of direct connections between the dura and the brain. Nature 627, 165–173 (2024).

52. Yoon JH, et al. Nasopharyngeal lymphatic plexus is a hub for cerebrospinal fluid drainage. Nature, (2024).

53. Haas J, et al. The Choroid Plexus Is Permissive for a Preactivated Antigen-Experienced Memory B-Cell Subset in Multiple Sclerosis. Front Immunol 11, 618544 (2020).

54. Liston A, Pasciuto E, Fitzgerald DC, Yshii L. Brain regulatory T cells. Nature Reviews Immunology, (2023).

55. Hauser SL, et al. Ocrelizumab versus Interferon Beta-1a in Relapsing Multiple Sclerosis. The New England journal of medicine 376, 221–234 (2017).

56. Montalban X, et al. Ocrelizumab versus Placebo in Primary Progressive Multiple Sclerosis. The New England journal of medicine 376, 209–220 (2017).

57. de Sèze J, et al. Anti-CD20 therapies in multiple sclerosis: From pathology to the clinic. Front Immunol 14, 1004795 (2023).

58. Wang T, et al. Unexpected BrdU inhibition on astrocyte-to-neuron conversion. Neural regeneration research 17, 1526–1534 (2022).

