## Supplemental Figures and Figure legends for "Antibodies against the capsid induced after intracranial AAV administration limits second administration in a dose dependent manner"

**Legends of supplemental figures**

**Figure S1 (related to Figure 1). The universally compromising effect of high-dose AAV on the second round AAV of the same serotype.**

(A) The representative images showing that the first (tdT in red) and second (GFP in green) round of AAV5 can expressed in same cells (in yellow) when using low-dose AAV5 in the first round (Low, top panel), while the expression of the second round of AAV5 is almost compromised when using high-dose AAV5 (High, bottom panel) in the first round. Scale bar: 20 μm

(B) The representative images revealing that the second round of AAV5 (GFP) is compromised by the middle-dose (1.2E10 vg/mouse) of intrastriatal injection of AAV5 as the first round (tdT in red). Scale bars: 200 μm

(C-D) Experimental design (C) and representative images (D) showing that the compromising effect of high-dose intrastriatal injection of AAV5 on the redosing is independent of transgenes. Note that switching the order of transgenes in the first and second round, the re-administration is also compromised. The corpus callosum and lateral ventricle are delineated by the white dashed lines. Scale bar: 200 μm

(E-F) Experimental design (E) and representative images (F) showing that increasing the dosage of second round of AAV from 2E9 vg to 4E10 vg fails to circumvent the compromising effect. The corpus callosum and lateral ventricle are delineated by the white dashed lines. Scale bar: 200 μm

(G-H) Experimental design (G) and representative images (H) showing that the compromising effect of high-dose intrastriatal injection of AAV on the redosing is independent of AAV serotype. Note that switching the serotype of AAV from AAV5 to AAV9, the re-administration is also compromised. The corpus callosum and lateral ventricle are delineated by the white dashed lines. Scale bar: 200 μm

(I-K) Quantitative analyses showing the fluorescence intensity of the second-round AAV-delivered gene after receiving the first-round intrastriatal injection of high-dose AAV is much lower than that receiving PBS injection, regardless of transgene (I), the dose of the second-round virus (J) and serotype (K). Mean (SD); unpaired t test, ***$P = 0.0003$ for J, ****$P < 0.0001$ for I and K, n= 3-5 mice/group

### Figure S1

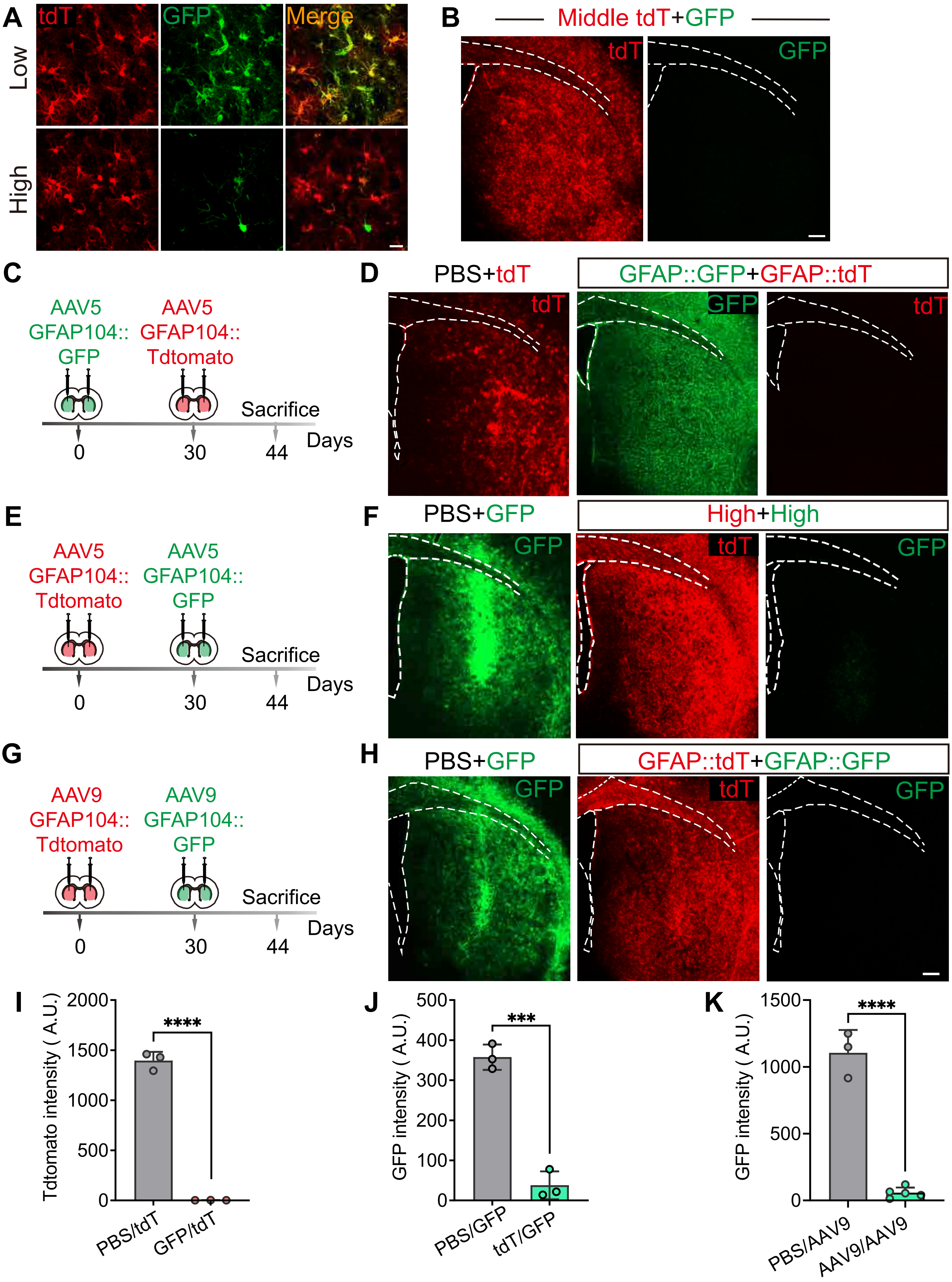

A
tdT
GFP
Merge
Low
High
B
Middle tdT+GFP
tdT
GFP
C
AAV5 GFAP104:: GFP
AAV5 GFAP104:: Tdtomato
Sacrifice
Days
0
30
44
D
PBS+tdT
GFAP::GFP+GFAP::tdT
tdT
GFP
tdT
E
AAV5 GFAP104:: Tdtomato
AAV5 GFAP104:: GFP
Sacrifice
Days
0
30
44
F
PBS+GFP
High+High
GFP
tdT
GFP
G
AAV9 GFAP104:: Tdtomato
AAV9 GFAP104:: GFP
Sacrifice
Days
0
30
44
H
PBS+GFP
GFAP::tdT+GFAP::GFP
GFP
tdT
GFP
I
Tdtomato intensity ( A.U.)
2000
1500
1000
500
0
****
PBS/tdT
GFP/tdT
J
GFP intensity ( A.U.)
500
400
300
200
100
0
***
PBS/GFP
tdT/GFP
K
GFP intensity ( A.U.)
1500
1000
500
0
****
PBS/AAV9
AAV9/AAV9

**Figure S2 (related to Figure 1). The universal compromising effect of high-dose AAV on the expression of second round AAV of the same serotype is independent of targeting cells.**

(A-B) Experimental design (A) and representative images (B) revealing that using high dose of AAV5 GFAP104::tdT (red) to target astrocytes in the first round compromises the expression of AAV5 Syn::GFP (green) to target neurons in the second round. The corpus callosum and lateral ventricle are delineated by the white dashed lines.

(C-D) Experimental design (C) and representative images (D) revealing that using high dose of AAV5 Syn::mCh (mCherry in red) to target neurons in the first round compromises the expression of AAV5 GFAP104::GFP (green) to target astrocytes in the second round. The corpus callosum and lateral ventricle are delineated by the white dashed lines. Scale bars: 200 μm

(E-F) Quantitative analysis showing the exponential decrease in the expression of second transgene (GFP) delivered by the first round of high-dose AAV5 injection, regardless of using astrocyte-specific promoter (E related to A and B) or neuron-specific promoter (F related to C and D). Mean (SD); unpaired t test, ***P= 0.0002 for E, **P= 0.0011 for F, n= 3-5 mice/group.

### Figure S2

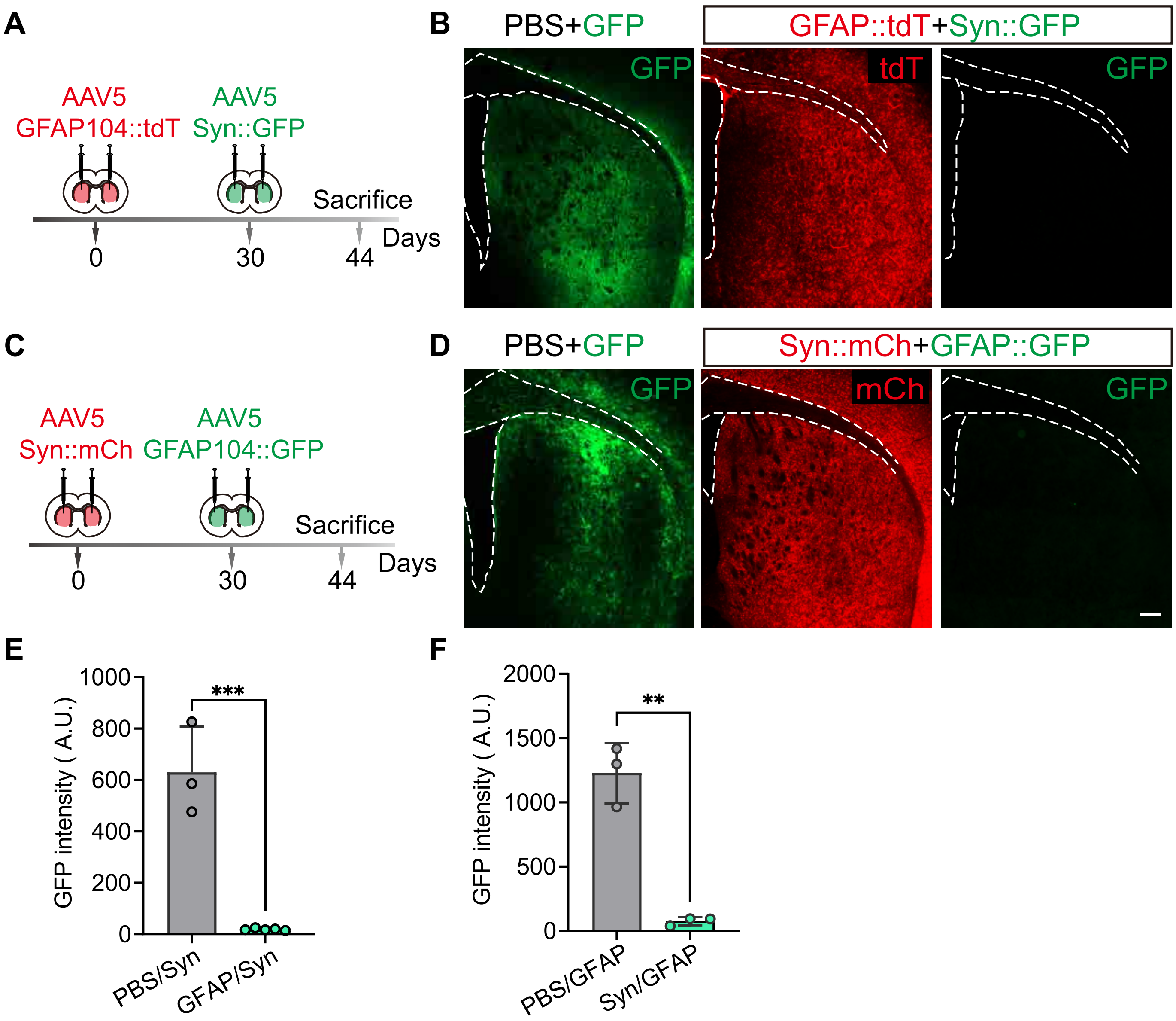

A
AAV5
GFP104::tdT
AAV5
Syn::GFP
Sacrifice
0
30
44
Days
B
PBS+GFP
GFAP::tdT+Syn::GFP
GFP
tdT
GFP
C
AAV5
Syn::mCh
AAV5
GFAP104::GFP
Sacrifice
0
30
44
Days
D
PBS+GFP
Syn::mCh+GFAP::GFP
GFP
mCh
GFP
E
GFP intensity ( A.U.)
1000
800
600
400
200
0
***
PBS/Syn
GFAP/Syn
F
GFP intensity ( A.U.)
2000
1500
1000
500
0
**
PBS/GFAP
Syn/GFAP

**Figure S3 (related to Figure 3) High-dose AAV induces immune response in both injection site and non-injection site**

(A) Experimental design to investigate the RNA profile of the injection region (striatum) and non-injection region (hippocampus) 21 days after performing intrastriatal microinjection of high-dose AAV5. PBS is used as control group. N= 3 mice/group

(B) Bar graph showing 58 DEGs (FDR< 0.05) in the hippocampus after intrastriatal injection.

(C) The KEGG-enriched bar graph showing that the pathways in the hippocampus induced by intrastriatal microinjection of high-dose AAV are mainly restricted to the immune response (one-way ANOVA analysis with Tukey test).

(D -E) Volcano map plotting the DEGs in striatum (D) and hippocampus (E) after high-dose AAV5 administration. Representative DEGs are marked in black, corresponding to blue dots.

(F-G) Representative images (F) and quantitation (G) showing that astrocytes (GFAP in green) in the striatum receiving high-dose AAV5 upregulates the expression of VIMENTIN (red) representing the reactive astrocytes. Scale bar: 200 μm; dpi: days post injection. Mean (SD); unpaired t-test, *P= 0.0254, n= 3 mice/group.

(H-I) Representative images (H) and quantitation (I) showing that increased signal of CD68 (green) in the striatum receiving high-dose AAV5. Scale bar: 20 μm. Mean ± SD; unpaired t-test, **P= 0.0012, n= 5 mice/group.

(J-K) Low-magnification images of the overall expression of CXCL10 (J) and CXCL9 (K) in injected area (related to Figure 3J and 3L). Scale bars: 200 μm

(L) Representative images revealing the localization and distribution of CXCL9 positive cells (red). Note that the location of the CXCL9 positive cells is closely related to blood vessel (PDGFRB in blue). A part of astrocytes (GFAP in green) locating adjacent to the blood vessel express CXCL9, while most of the CXCL9 positive cells locate in the space between blood vessel and astrocytic limitans. Scale bars: 20 μm

(M) Typical images showing that in the hippocampus which is the non-injection region of the first round of AAV5, the signal of GFAP, IBA1, CD45, CXCL9 and CXCL10 is comparable to that in the PBS group. The cell nuclei are counterstained with DAPI (blue). Scale bars: 500 μm

### Figure S3

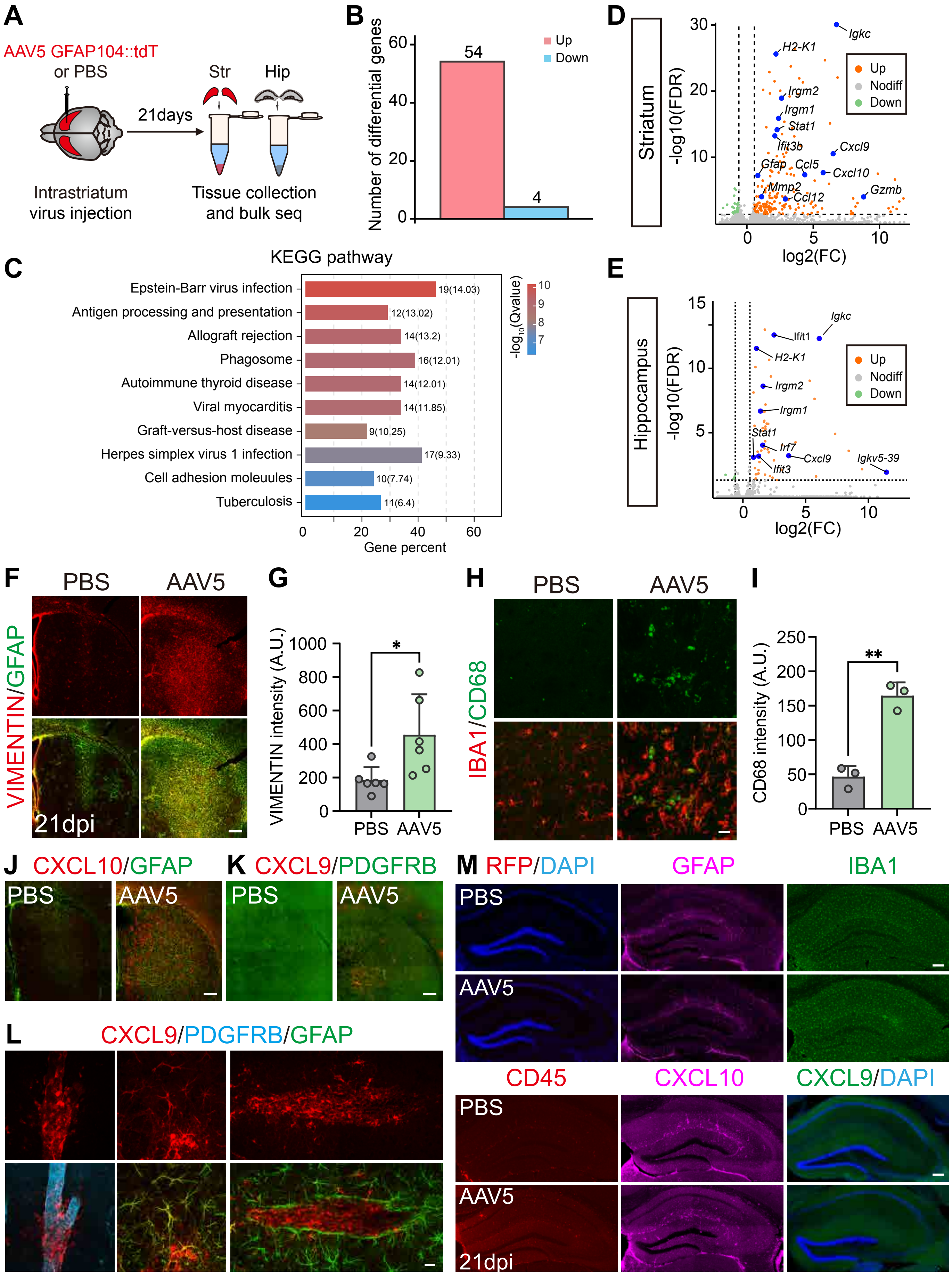

A
AAV5 GFAP104::tdT
or PBS
21days
Str
Hip
Intrastriatum
virus injection
Tissue collection
and bulk seq
B
Number of differential genes
54
4
Up
Down
C
KEGG pathway
Epstein-Barr virus infection
Antigen processing and presentation
Allograft rejection
Phagosome
Autoimmune thyroid disease
Viral myocarditis
Graft-versus-host disease
Herpes simplex virus 1 infection
Cell adhesion moleuules
Tuberculosis
19(14.03)
12(13.02)
14(13.2)
16(12.01)
14(12.01)
14(11.85)
9(10.25)
17(9.33)
10(7.74)
11(6.4)
Gene percent
D
Striatum
-log10(FDR)
log2(FC)
Igkc
H2-K1
Irgm2
Irgm1
Stat1
Ifit3b
Cxcl9
Gfap
Ccl5
Cxcl10
Gzmb
Mmp2
Ccl12
Up
Nodiff
Down
E
Hippocampus
Igkc
Ifit1
H2-K1
Irgm2
Irgm1
Stat1
Irf7
Cxcl9
Igkv5-39
Ifit3
F
PBS
AAV5
VIMENTIN/GFAP
21dpi
G
VIMENTIN intensity (A.U.)
*
H
PBS
AAV5
IBA1/CD68
I
CD68 intensity (A.U.)
**
J
CXCL10/GFAP
K
CXCL9/PDGFRB
L
CXCL9/PDGFRB/GFAP
M
RFP/DAPI
GFAP
IBA1
CD45
CXCL10
CXCL9/DAPI
21dpi

**Figure S4 (related to Figure 4) High-dose AAV triggers T cell infiltration that may not the direct reason for the compromising effect**

(A) Representative images showing the infiltration of B220$^+$ cells in the hippocampus 21 days after intrastriatal injection of high-dose AAV5. Scale bar: 20 μm; dpi: days post injection.

(B) Typical picture showing that the spleen of B6-Ighj-KO mouse (KO) is smaller than that of the C57BL/6J wildtype mouse (C57).

(C) Representative images showing that B6-Ighj-KO mice (KO) have far fewer B220$^+$ cells (red) in the spleen than C57BL/6J mice (WT), while CD8$^+$ (green) and CD4$^+$ cells (magenta) are comparable. Scale bar: 200 μm

(D-E) Representative images revealing the robust invasion of T lymphocytes (CD4$^+$ and CD8$^+$ cells in red) in the striatum (D) and hippocampus (E) at 21 days after the high-dose AAV5 injection. The cell nuclei are counterstained with DAPI (blue). Scale bars: 20 μm, dpi: days post infection

(F) Quantification showing increased density of CD8$^+$ and CD4$^+$ cells in the striatum of wildtype mouse that received high-dose AAV5 administration. Mean (SD); two-way ANOVA with Sidak's multiple comparison test, *P= 0.0348 for CD4$^+$ cells and *P= 0.0475 for CD8$^+$ cells, n= 5 mice/group.

(G) Representative images showing that high-dose AAV5 triggered T lymphocytes (CD8$^+$ and CD4$^+$ cells in red) infiltration in the striatum of B6-Ighj-KO mouse. The cell nuclei are counterstained with DAPI (blue). Scale bar: 20 μm

(H) Quantification showing increased density of $CD8^+$ and $CD4^+$ cells in the striatum of B6-Ighj-KO mouse that received high-dose AAV5 administration. Mean (SD); two-way ANOVA with Sidak's multiple comparison test, **P= 0.0029 for $CD4^+$ cells and **P= 0.0018 for $CD8^+$ cells, n= 5 mice/group.

### Figure S4

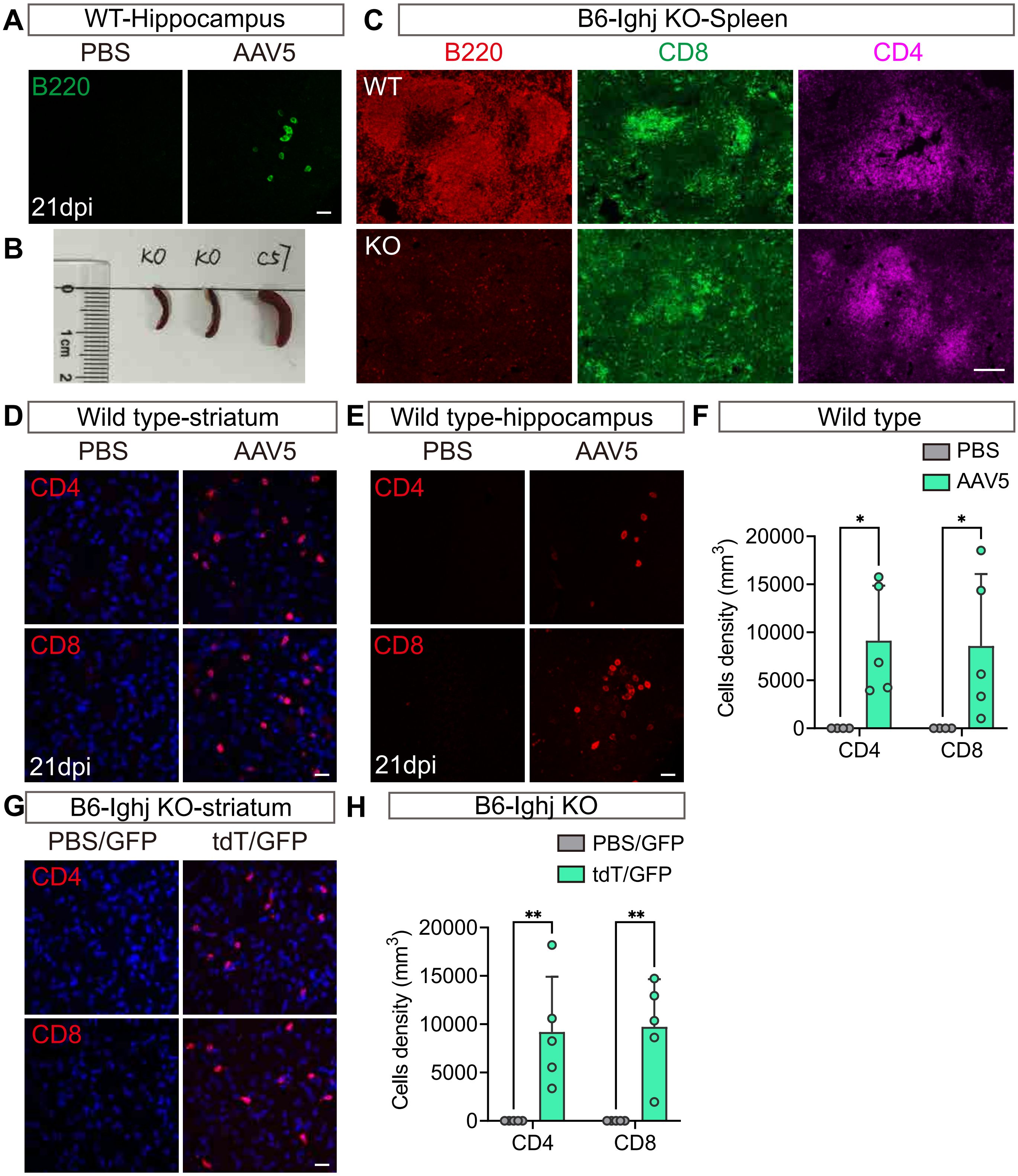

A
WT-Hippocampus
PBS
AAV5
B220
21dpi
B
KO
KO
C57
1 cm
2
C
B6-Ighj KO-Spleen
B220
CD8
CD4
WT
KO
D
Wild type-striatum
PBS
AAV5
CD4
CD8
21dpi
E
Wild type-hippocampus
PBS
AAV5
CD4
CD8
21dpi
F
Wild type
PBS
AAV5
Cells density (mm3)
20000
15000
10000
5000
0
*
*
CD4
CD8
G
B6-Ighj KO-striatum
PBS/GFP
tdT/GFP
CD4
CD8
H
B6-Ighj KO
PBS/GFP
tdT/GFP
Cells density (mm3)
20000
15000
10000
5000
0
**
**
CD4
CD8

**Figure S5 (related to Figure 5) Parenchymal administration of high-dose AAV5 did not disrupt BBB integrity**

(A) Representative images of the endothelial cells of blood vessels (indicated by CD31, green) in the striatum at different time points after receiving the intrastriatal injection of high-dose AAV5. The striatum that received PBS and was collected at 21 dpi is set as the control. Scale bars: 200 μm; dpi: days post injection.

(B) Quantitation revealing that the intensity of CD31 signal in the striatum is comparable among the mice receiving PBS and receiving high-dose AAV5. Mean (SD); one-way ANOVA with Dunnett's multiple comparisons, n= 3 mice/group.

(C) Typical images showing that the distribution of tight junction protein (indicated by ZO-1, green) on the endothelial cells (indicated by CD31, red) in the striatum at different time points after receiving the intrastriatal injection of high-dose AAV5. The striatum that received PBS and was collected at 21 dpi is set as the control. Scale bars: 20 μm.

(D) Quantification showing that the localization ratio of ZO-1signal on the $CD31^+$ blood vessels is comparable among the mice receiving PBS and receiving high-dose AAV5. Mean (SD); one-way ANOVA with Dunnett's multiple comparisons, n= 3 mice/group.

(E)Typical images showing that the distribution of pericytes (indicated by PDGFRB, green) in the striatum at different time points after receiving the intrastriatal injection of high-dose AAV5. The striatum that received PBS and was collected at 21 dpi is set as the control. Scale bar: 20 μm.

(F) Quantification showing that the density of PDGFRB$^{+}$ pericytes is comparable among the mice receiving PBS or receiving high-dose AAV5. Mean (SD); one-way ANOVA with Dunnett's multiple comparisons, n= 3 mice/group.

(G) Representative images showing that the distribution of AQP4 (labeled astrocyte endfeet, green) in the striatum at different time points after receiving the intrastriatal injection of high-dose AAV5. The striatum that received PBS and was collected at 21 dpi is set as the control. Scale bar: 200 μm and 20 μm for inlets.

(H) Quantification revealing that the intensity of AQP4 is comparable among that receiving PBS or receiving high-dose AAV5. Mean (SD); one-way ANOVA with Dunnett's multiple comparisons, n = 3 mice/group.

(I-K) Quantitative analysis further showing that the distribution of AQP4 were similar in PBS- and AAV5-treated mice at different time points. (I, 7dpi; J, 14dpi; K, 21dpi). The distribution of AQP4 is evaluated by the intensity of AQP4 signal of the indicated pixels along the white dashed lines in the inlets in G.

(L) Experimental design to assess of BBB permeability with tracer FITC-d20. FITC-d20 was injected through the retro-orbital venous sinus at 21dpi of AAV/PBS injection, and circulated for 2 h. Then mice were sacrificed for immunofluorescence.

(M) Representative images of the distribution of 20 kDa dextran (green) in liver (top panel) and brain (bottom panel). Note that FITC fluorescence signal is detected in liver but absent in brain parenchyma both in mice receiving intracranial injection of PBS or high-dose AAV5 mice. Normal mice that received PBS via retro-orbital venous sinus served as negative control. The cell nuclei are counterstained with DAPI (blue). Scale bar: 20 μm

(N) Quantifications of the intensity of relative FITC fluorescence. Mean (SD); two-way ANOVA with Sidak's multiple comparison test, *P= 0.0189, n= 3 mice/group.

### Figure S5

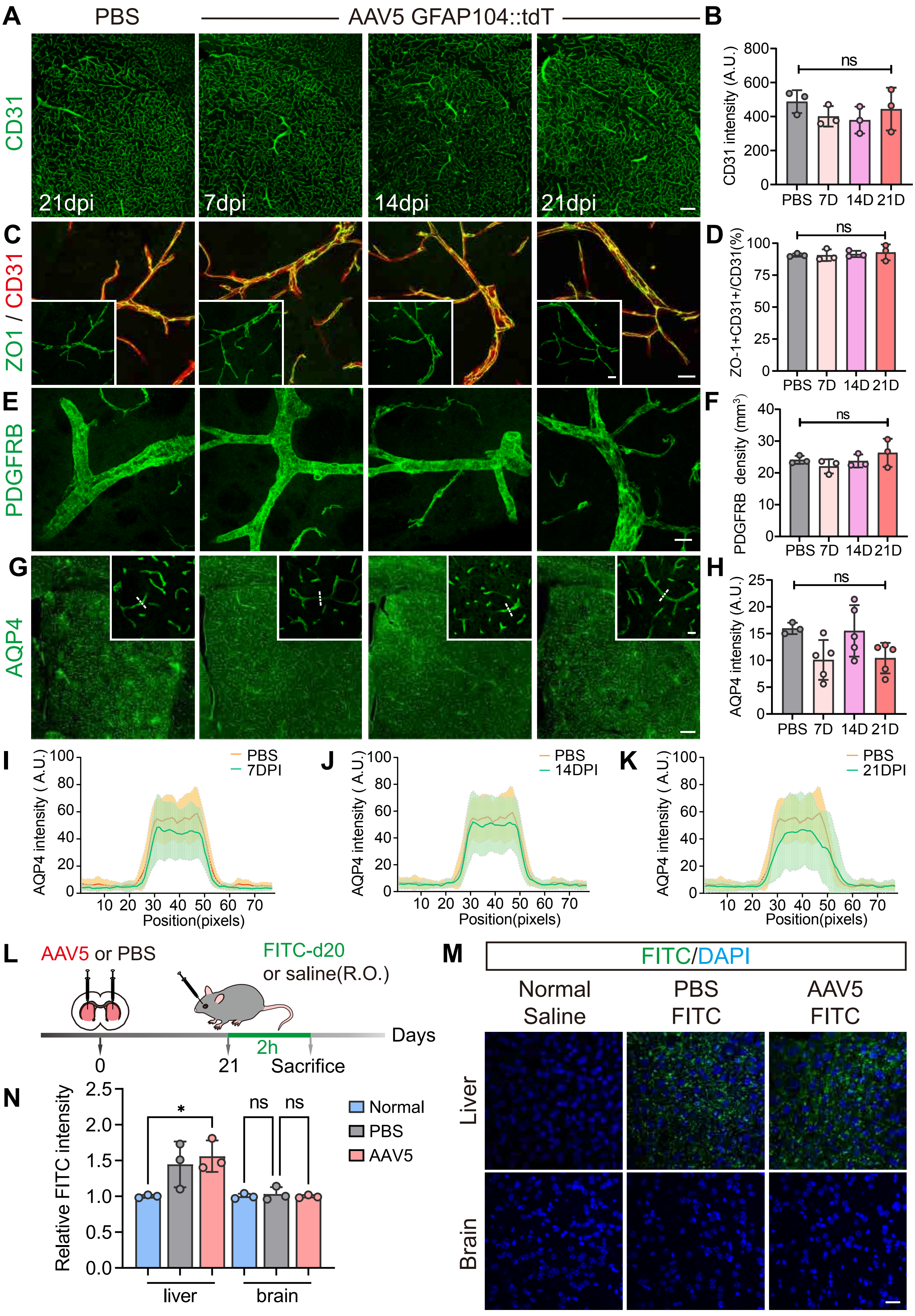
A
PBS
AAV5 GFAP104::tdT
CD31
21dpi
7dpi
14dpi
21dpi
B
CD31 intensity (A.U.)
ns
PBS 7D 14D 21D
C
ZO1 / CD31
D
ZO-1+CD31+/CD31(%)
E
PDGFRB
F
PDGFRB density (mm³)
G
AQP4
H
AQP4 intensity (A.U.)
I
PBS
7DPI
J
14DPI
K
21DPI
AQP4 intensity ( A.U.)
Position(pixels)
L
AAV5 or PBS
FITC-d20
or saline(R.O.)
Days
2h
0
21
Sacrifice
M
FITC/DAPI
Normal Saline
PBS FITC
AAV5 FITC
Liver
Brain
N
Relative FITC intensity
Normal
PBS
AAV5
liver
brain

**Figure S6 (related to Figure 5) Increased permeability of ventricular system nearby the high-dose AAV injection sites**

(A) Experimental design to assess the clearance of OA647 injected in the ventricular system. OA647 was injected in the right lateral ventricle at 21 dpi after AAV/PBS injection. 120 minutes later, the mice were sacrificed for immunofluorescence.

(B) Typical images of serial sections in the coronal plane showing that 120 minutes after OA647 (magenta) injection, the tracer retained in the brain parenchyma is comparable between that receiving PBS (top panel) and high-dose AAV5 (bottom panel). c', c'', d', and d'' are high-magnification images of areas delineated by the white dashed square in the low magnification images. The cell nuclei are counterstained with DAPI (blue). Scale bars: 1000 μm and 20 μm for c', c'', d', and d''; A: anterior; P: posterior; dpi: days post infection.

(C) Quantitation showing that the intensity of OA647 on the ipsilateral side is comparable between the mice receiving PBS and high-dose AAV5 120 minutes after dye injection. Mean (SD); unpaired t-test, n= 4 mice/group.

(D) Quantitative analysis showing the ratio of the intensity of OA647 on the ipsilateral side (where the dye was injected) to that on the contralateral side in the AAV5 group were greater than that in the PBS group when checked at 35 minutes after dye injection. But when checked at 120 minutes after the dye injection, the ratio is comparable. Mean (SD); two-way ANOVA with Sidak's multiple comparison test, *P= 0.0167, n= 4 mice/group.

(E) Representative images showing that the intensity of CXCL10 (red) increases in the regions around the lateral ventricle of the mice receiving high-dose AAV5 (right) compared with that receiving PBS (left) at 21 dpi. Str: striatum; LV: lateral ventricle: dpi: days post injection; scale bar: 20 μm.

(F) Representative images showing that the astrocytes (indicated by GFAP, green) nearby the lateral ventricle elevate the expression of CXCL10 (red) in the striatum of the mice receiving high-dose AAV5 (bottom panel) compared with that receiving PBS (top panel) at 21 dpi. The lateral ventricle regions are delineated by the white dashed lines. Str: striatum; LV: lateral ventricle: dpi: days post injection; scale bar: 20 μm.

### Figure S6

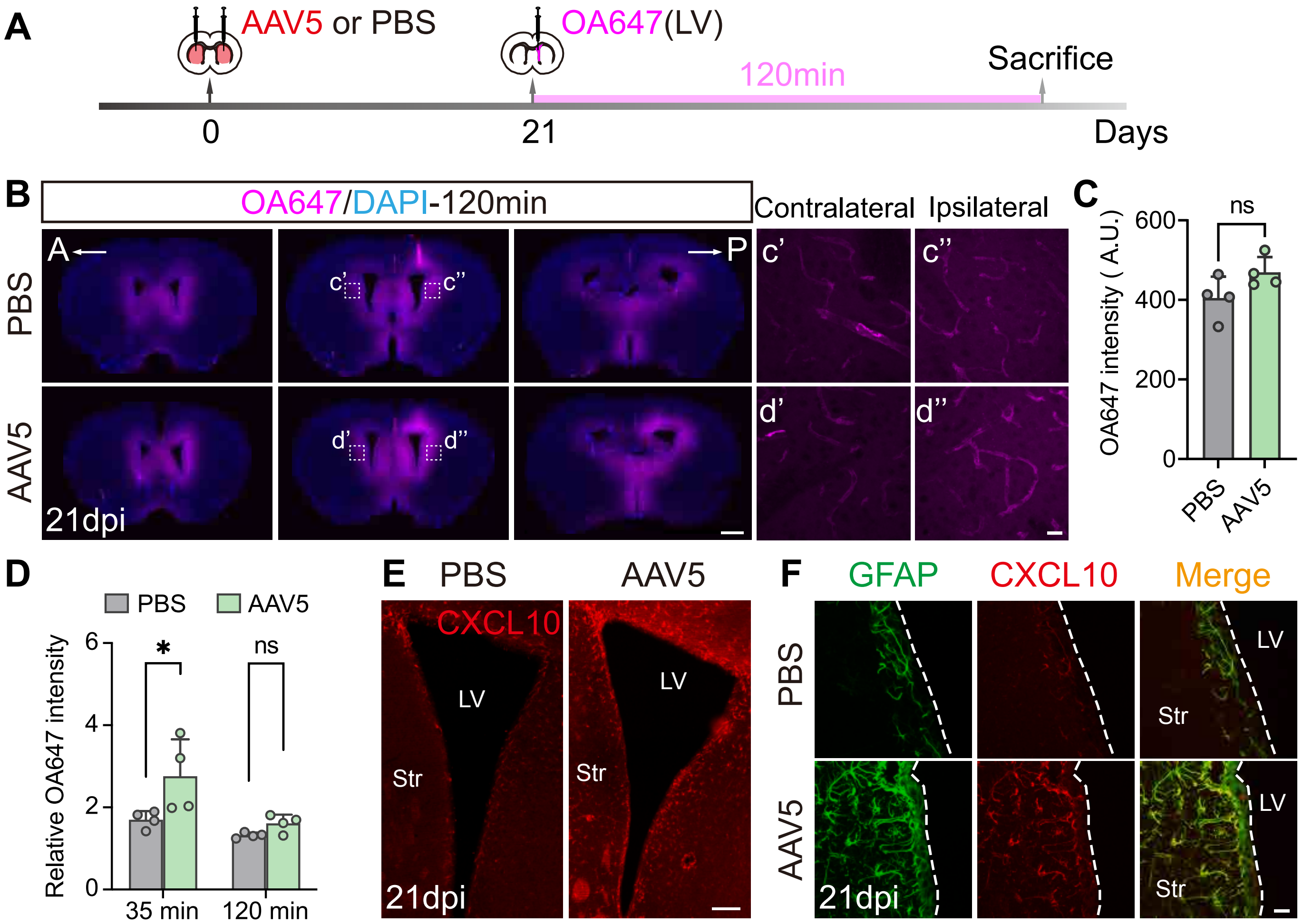
A
AAV5 or PBS
OA647(LV)
120min
Sacrifice
0
21
Days
B
OA647/DAPI-120min
Contralateral
Ipsilateral
A
P
PBS
AAV5
21dpi
c'
c''
d'
d''
C
ns
OA647 intensity ( A.U.)
600
400
200
0
PBS
AAV5
D
PBS
AAV5
Relative OA647 intensity
*
ns
6
4
2
0
35 min
120 min
E
PBS
AAV5
CXCL10
LV
Str
21dpi
F
GFAP
CXCL10
Merge
PBS
AAV5
LV
Str
21dpi

**Figure S7 (related to Figure 6) The treatment of AAV empty, IdeZ and HDAC3 inhibitor failed to circumvent the immunosuppression of NAbs**

(A) Experimental design. Mice that received high-dose AAV5 GFAP104::tdTomato in the first round were divided into four groups at 29 dpi. Two groups of mice were injected with AAV5 empty virus (1E11 GC/animal) via the retro-orbital sinus or lateral ventricle and one group was injected with PBS. One day later (at 30 dpi of the first round of AAV injection), AAV5 GFAP104::GFP was injected in the same location as the first round. In 4$^{th}$ group, the AAV5 GFAP104::GFP was combined with AAV5 empty virus before the intrastriatal injection. All groups were sacrificed at 14 dpi after receiving AAV5 GFAP104::GFP.

(B) Typical images showing the expression of the first round AAV5 (RFP, red) and the second round of AAV5 (GFP, green). Note the absence of GFP signal (bottom panel) suggesting that the treatment with AAV5 empty virus fails to improve the second transgene expression. Scale bar: 200 μm

(C) Experimental design of IdeZ treatment. Lateral ventricle administration of IdeZ (an IgG-degrading enzyme) to transiently cleave parenchymal NAbs, which was performed one day before the second round of virus injection. Mice administrated with PBS serve as controls.

(D) Representative images showing the absence of GFP (green) signal indicating the expression of second-round AAV5 in PBS (left) and IdeZ (right) treated mice. Note that the expression of the first-round AAV5 (indicated by RFP, red) is normal. Scale bar: 200 μm

(E) Experimental design of RGFP966 treatment. Mice firstly received high-dose AAV5 GFAP104::tdTomato, and then high-dose AAV5 GFAP104::GFP at a 21-day interval. From 5 to 24 days after the first injection, the mice were administrated either DMSO or RGFP966 (a HDAC3 inhibitor) by intraperitoneal injection once a day.

(F) Representative images showing that the GFP signal indicating the expression of second-round AAV5 is absent in DMSO and RGFP966 treated group. Note that the expression of the first-round AAV5 (indicated by RFP, red) is normal. Scale bar: 200 μm

(G-I) Quantifications of GFP intensity after AAV5 empty (G), IdeZ (H) and RGDP966 (I) treatment. Mean (SD); one-way ANOVA with Dunnett's multiple comparisons or unpaired t test, n= 3-5 mice/group.

### Figure S7

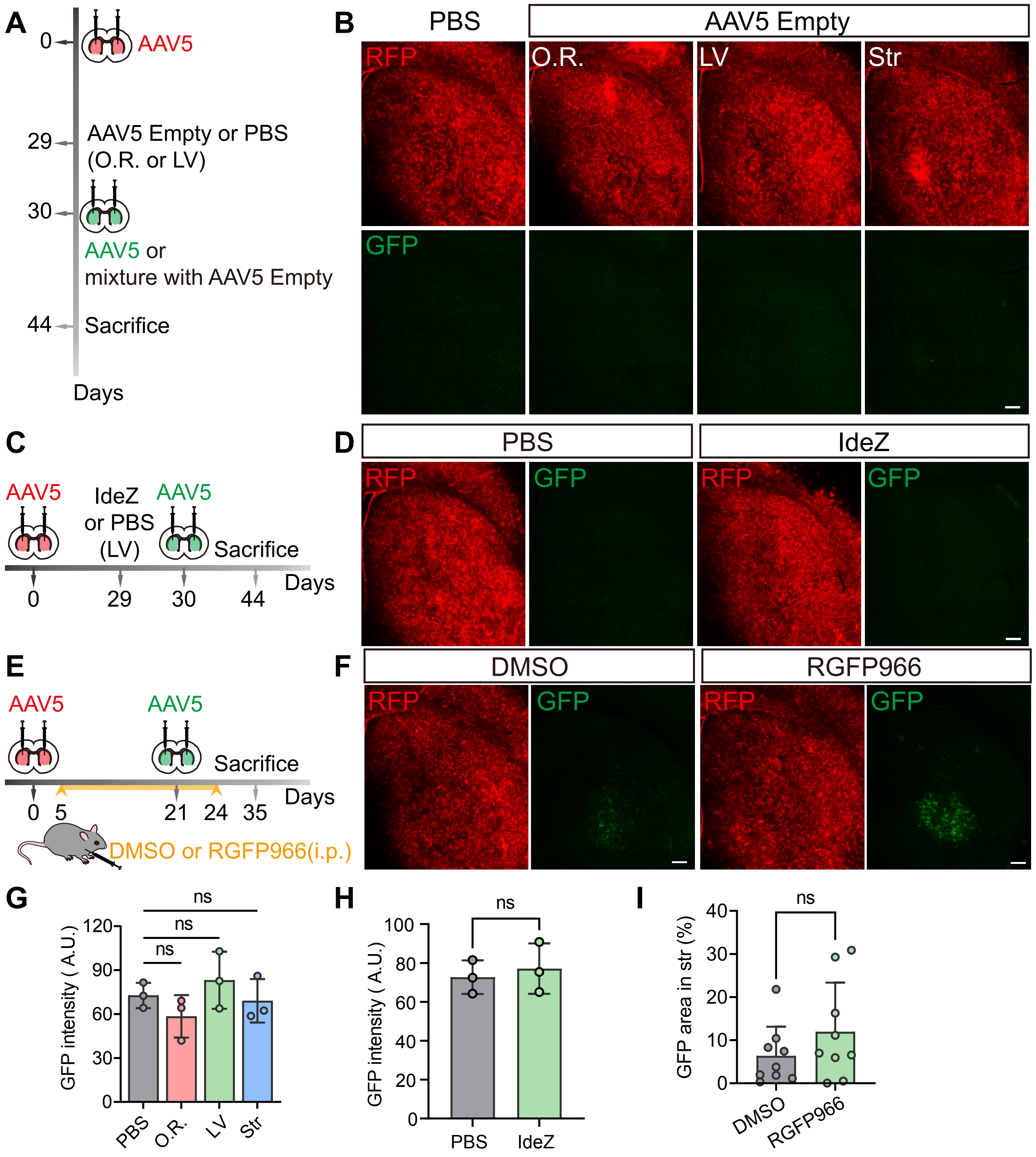
A
0
AAV5
29
AAV5 Empty or PBS
(O.R. or LV)
30
AAV5 or
mixture with AAV5 Empty
44
Sacrifice
Days
B
PBS
AAV5 Empty
RFP
O.R.
LV
Str
GFP
C
AAV5
IdeZ
or PBS
(LV)
AAV5
Sacrifice
Days
0
29
30
44
D
PBS
IdeZ
RFP
GFP
RFP
GFP
E
AAV5
AAV5
Sacrifice
Days
0
5
21
24
35
DMSO or RGFP966(i.p.)
F
DMSO
RGFP966
RFP
GFP
RFP
GFP
G
ns
ns
ns
GFP intensity ( A.U.)
120
90
60
30
0
PBS
O.R.
LV
Str
H
ns
GFP intensity ( A.U.)
100
80
60
40
20
0
PBS
IdeZ
I
ns
GFP area in str (%)
40
30
20
10
0
DMSO
RGFP966

**Table S1 Primary antibody information**

| Antibody | Host | Dilution | Source | Catalog |
|---|---|---|---|---|
| AQP4 | Rabbit | 1:1000 | Proteintech | 16473-1-AP |
| B220 | Rat | 1:200 | Invitrogen | 14-0452-82 |
| CD31 | Goat | 1:200 | Biotechne | AF3628 |
| CD45 | Rat | 1:1000 | BD Bioscience | 550539 |
| CD4 | Rat | 1:500 | ThermoFisher | 14-0042-82 |
| CD68 | Rabbit | 1:500 | Abcam | Ab125212 |
| CD8 | Rabbit | 1:500 | Abcam | Ab217344 |
| CD138 | Rabbit | 1:500 | Proteintech | 15093-1-AP |
| CXCL9 | Goat | 1:200 | BD Systems | AF-492-NA |
| CXCL10 | Goat | 1:200 | BD Systems | AF-466-NA |
| GFAP | Rat | 1:1000 | Invitrogen | 13-0300 |
| IBA1 | Rabbit | 1:500 | Wako | 019-19741 |
| PDGFRB | Goat | 1:200 | BD Systems | AF1042 |
| PDGFRB | Rat | 1:200 | Invitrogen | 14-1402-82 |
| ZO-1 | Rat | 1:200 | DSHB | R26.4C |
